# Investigating Overflow Metabolism in Heterotrophic Cultures of the Green Alga *Chromochloris zofingiensis*

**DOI:** 10.1101/2025.10.20.683564

**Authors:** Michelle Meagher, Dimitrios J. Camacho, Sean D. Gallaher, Sabeeha S. Merchant, Nanette R. Boyle

## Abstract

*Chromochloris zofingiensis* is of interest for its ability to perform a reversible trophic switch in the presence of glucose that is characterized by a shutdown of photosynthesis and an accumulation of energy storage metabolites. Previous work has shown that this trophic switch is accompanied by overflow metabolism and the production of lactate in aerobic conditions. This trophic switch is not observed in nutrient replete media. We utilized isotopically assisted metabolic flux analysis to characterize intracellular flux distributions that are associated with different metabolic phenotypes observed in this organism in different media formulations in light and dark conditions. The results of this analysis showed differences in flux through carbon fixation reactions, the TCA cycle and through the reaction catalyzed by pyruvate kinase. This analysis was complemented with transcriptomics data collected for *C. zofingiensis* grown in iron limited conditions to provide further evidence towards the negative impact of iron limitation on both photosynthetic and respiratory activity. Overflow metabolism allows this alga to compensate for the lower energy production that results from iron limitation. This work highlights how nutrient availability can lead to drastic changes in the metabolism of *C. zofingiensis*.

## Introduction

*Chromochloris zofingiensis* is an emerging model green alga which is of interest for its ability to produce high levels of triacylglycerol [1] in addition to the production of astaxanthin, a high value pigment with powerful antioxidant capabilities. During growth on glucose, *C. zofingiensis* undergoes a trophic switch from autotrophy to heterotrophy, even under continuous illumination [2]. This reversible trophic switch is characterized by an accumulation of lipids, starch, and astaxanthin within the cell, alongside the downregulation of chlorophyll production and loss of the photosynthetic machinery [2]. This loss can be avoided if sufficient iron is provided to the culture [3], allowing for mixotrophic growth on glucose.

Iron is an essential micronutrient that plays a critical role in photosynthetic and respiratory electron transport chains, and is a necessary cofactor for many enzymes catalyzing metabolic reactions. In algae, iron limitation can inhibit photosynthetic energy generation [4-6]. Plants respond in a similar manner to iron limitation, and in addition to photosynthesis [7], respiration is impacted in root tissue [8]. In heterotrophically growing organisms like yeast, iron deficiency limits respiration and when fermentative substrate is available, these organisms reprioritize limited iron towards other necessary cell processes shifting metabolism to fermentation for energy generation [9].

In previous work, a genome scale metabolic model, iCzof1925, was created for *C. zofingiensis*. Flux balance analysis (FBA) performed using this model accurately predicted the fermentation of glucose during heterotrophic growth in *C. zofingiensis,* with lactic acid as the primary fermentation product excreted [10]. In this work, we show that when *C. zofingiensis* is grown in a nutrient optimized medium [11] containing glucose, lactic acid excretion does not occur, and photosynthetic capacity of the cells is retained. To further investigate this phenomenon, we conducted an isotopically assisted metabolic flux analysis (MFA) of heterotrophic *C. zofingiensis* cultures grown on glucose in both a basic and a nutrient replete media. The results of this analysis indicated that *C. zofingiensis* cultures grown in a standard medium commonly used for green algae had an altered metabolism as a result of nutrient limitation, with iron limitation identified as a major contribution to this phenotype. To further support these findings, transcriptomics data collected from *C. zofingiensis* in three different iron concentrations is presented. Changes in gene expression for various enzymes are compared to MFA flux data for pathways containing these enzymes to provide a more detailed description of the metabolic changes that occur in *C. zofingiensis* during growth in suboptimal nutrient conditions.

## Methods

### Metabolic Flux Analysis Experiments

#### Strains and Culturing Conditions

*Chromochloris zofingiensis* (strain ID SAG 211-14) cultures were generously provided by the Niyogi Laboratory at UC Berkeley. The nutrient replete medium type was developed by collaborators at UC Berkeley and has been designated as CORE medium (Chromochloris Optimized Ratio of Elements) [11]. The other medium formulation used was Tris-phosphate media, a variation on TAP medium [12] with glucose replacing acetate as an organic carbon source, named TGP in work presented here. This TGP medium used Hutner’s trace element solution purchased from the Chlamydomonas Resource Center at a final concentration of 1 mL/L. Filter sterilized glucose was added to both media types to a final concentration of 20 g/L. Cultures were grown in 250 mL baffled shake flasks agitated at a rate of 180 rpm in incubators held at 25 °C, and for continuous light experiments, light intensity was 40 µmol/m^2^/s supplied using Phillips F15T8 bulbs (2 4100K : 1 3000K).

### Spent Media Analysis

Growth was monitored for all growth conditions, during which time culture samples were collected daily, filtered with 0.2 µm pore size filters, and the supernatant was frozen for spent media analysis. Spent media samples were analyzed on the YSI 2950 biochemistry analyzer to quantify glucose uptake. To quantify fermentation product secretion, dried spent media samples were derivatized and analyzed on GC-MS as described by Young et.al. [13], and identified peaks were quantified against a ribitol internal standard. Iron concentrations in fresh media and filtered media collected during mid-exponential growth phase were determined using a Cedex Bioanalyzer ferrozine based iron assay.

### Isotope Labeling Experimental Design

When conducting isotopically assisted metabolic flux analysis, labeling experiments must be carefully designed to maximize information useful for flux determinations while also minimizing the cost of expensive isotopically labeled reagents [14]. Both the selection of tracers and the structure of the metabolic network have a large impact on the predicted flux distributions [14]. The glucose tracer 1,2-^13^C glucose has been found to be optimal for flux resolution in the pentose phosphate pathway and glycolysis [15, 16], while U-^13^C glucose has been found to provide good flux resolution for the TCA cycle [16], so these labeled substrates were used as the focus for isotopic tracer simulations. Isotopomer Network Compartmental Analysis (INCA) [17] software enables the user to conduct isotopic tracer simulations, to evaluate anticipated labeling patterns *in silico*, and evaluate their usefulness in the determination of fluxes. INCA was used to conduct isotopic tracer simulations on the central metabolic network of *C. zofingiensis*, testing various combinations of unlabeled [U-^12^C], uniformly labeled [U-^13^C], and [1,2-^13^C] labeled glucose. Based on the results of these simulations, a 60:20:20 mixture of [U-^12^C]: [U-^13^C]: [1,2-^13^C] glucose was selected for this study.

Prior to the inoculation of experimental cultures, seed flasks of *C. zofingiensis* in both media types were grown on isotopically labeled glucose for 5 days. This was done to ensure that the cultures reach isotopic steady state (i.e. the isotopic labeling has fully penetrated throughout the metabolic network). After this point, seed cultures were used to inoculate experimental cultures which were grown to mid exponential phase to ensure metabolic steady state before harvesting, quenching, and storing the frozen samples until further analysis. Quenched, labeled cell pellets were analyzed to obtain mass isotopomer distributions for intracellular metabolites listed in supplemental table S1.

Figure **1** below shows an outline of the workflow for these experiments.

**Figure 1.**
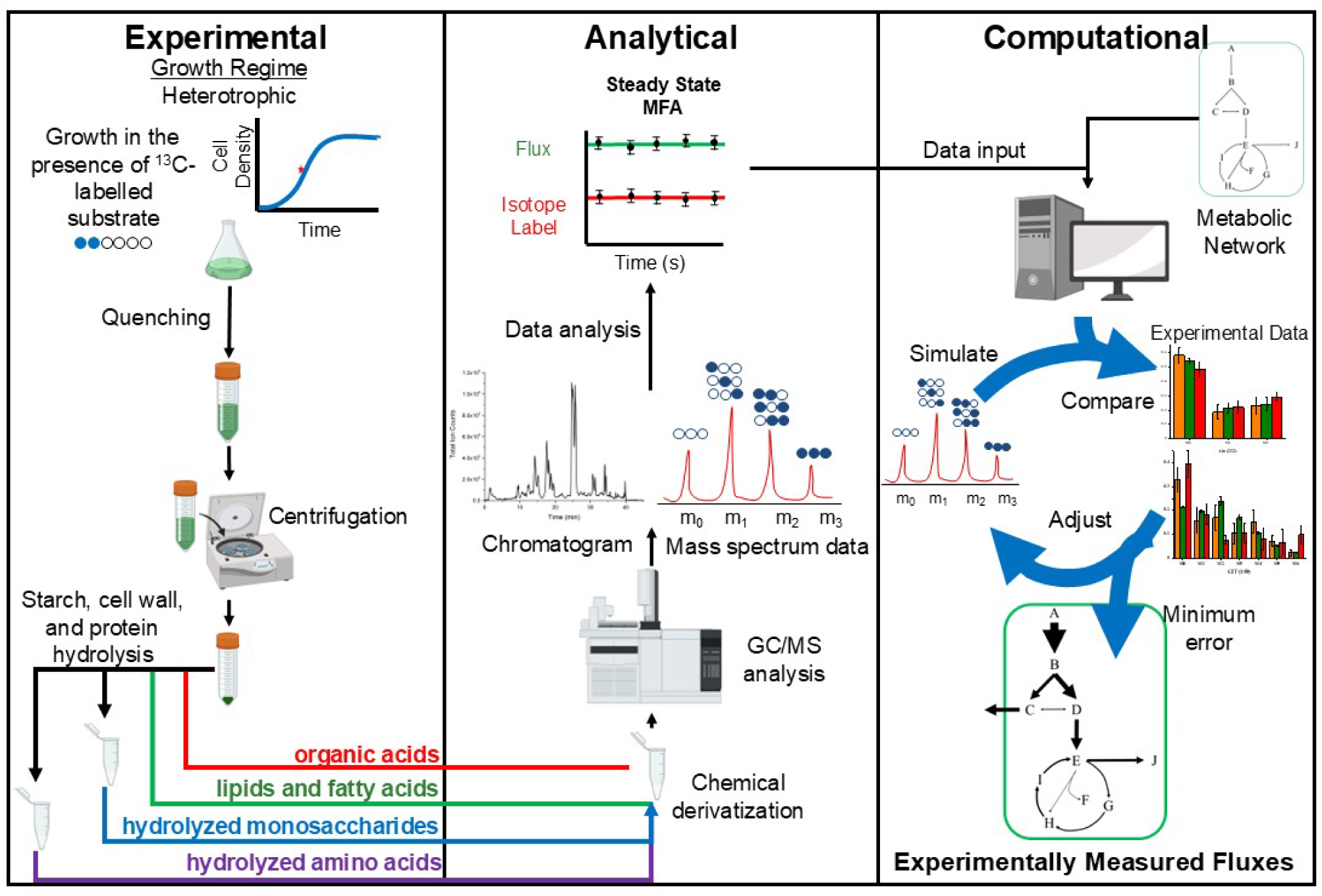
Workflow diagram for ^13^C-MFA in heterotrophic cultures. Heterotrophic algae cultures are grown in the presence of ^13^C labeled glucose until isotopic steady state is achieved and the ^13^C label has penetrated throughout the metabolic network. Cultures are then harvested during mid-exponential phase while at metabolic steady state. Upon harvesting, samples are quenched to rapidly halt metabolism, and quenched cell pellets are processed to derivatize and analyze amino acids, sugars, fatty acids, and organic acids via GC-MS. Mass isotopomer distributions from this analysis are used along with external flux data and a central metabolic network model to estimate fluxes through the network. This is done in an iterative process, adjusting flux values to minimize error between simulated and measured isotopomer distributions.

### Sample Collection and Quenching

Culture samples were quenched upon collection to rapidly stop metabolic activity and preserve *in vivo* metabolite concentrations and isotopic labeling patterns. The quenching procedure was carried out as described by Sake et.al. [18], and the quenched cell pellets were stored at −28°C until analysis.

### Amino Acids Sample Preparation

For analysis of isotopic labeling of amino acids, proteins were hydrolyzed in 6 M HCl under a vacuum for 20 h, and the resulting mixture of amino acids was converted to their tert-butyldimethylsilyl derivatives as described in work by Antoniewicz et. al. [19]. After derivatization, the mixture was analyzed via GC-MS. The localization of amino acid biosynthesis reactions in the genome-scale metabolic model of *C. zofingiensis* [10] were used to predict the compartments in which each amino acid is synthesized, allowing for compartment specific labeling data from amino acids.

### Organic Acids Sample Preparation

Organic acids were extracted from quenched cell pellets prior to analysis. Metabolite extraction was performed using the extraction procedure described by Young et.al. [20], modified to incorporate grinding in a cryogenic ball mill to lyse the thick cell walls present in *C. zofingiensis*. Quenched cell pellets were ground with 5 mm diameter stainless steel balls in a Retch cryomill under liquid nitrogen cooling for 2.5 min at 25 Hz. After grinding, 500 μL of Optima grade methanol was added, and samples were resuspended by pipetting up and down before being transferred to a fresh tube. Samples were then subjected to three freeze-thaw cycles consisting of dipping in liquid nitrogen and thawing on a shake plate at 4 °C before collecting the supernatant after centrifugation at 10,000 ×*g* at 1 °C for 5 minutes. The supernatant was transferred to another fresh tube. The remaining pellet was extracted twice more as before using a 50:50 mixture of Optima water and methanol. Supernatants from each extraction procedure were pooled and dried on a Thermo Fisher SpeedVac Concentrator. Dried samples were then derivatized and analyzed via GC-MS [13].

### Carbohydrates sample preparation

Cell wall sugars, produced in the cytosol, and starch, produced in the plastid, were analyzed to gain compartment specific labeling information on glucose in the cytosol and plastid. To analyze cell wall sugars, biomass was hydrolyzed with HCl as described by McConnell and Antoniewicz [21]. Cellular starch was hydrolyzed as described by Young et.al. [13]. Dried samples containing sugar monomers from the above procedures were derivatized with hydroxylamine hydrochloride (2% w/v) in pyridine and propionic anhydride [13]. After derivatization, the mixture was analyzed via GC-MS.

### Fatty Acids Analysis

To obtain compartment specific labeling information for acetyl-CoA in the chloroplast, fatty acids were analyzed via GC-MS. Fatty acids from quenched cell pellets were converted to their fatty acid methyl ester (FAME) derivatives and analyzed as described by Christie et.al. [22].

### GC-MS Analysis

All work was performed on an Agilent 6890 gas chromatograph with an Agilent 5973 single quadrupole mass spectrometer detector using helium as the carrier gas. Amino acids, organic acids, and sugars were analyzed on a DB-1701 column from Agilent, and fatty acids were analyzed on a DB-WAX column from Agilent. GC-MS operating conditions were as described in the literature for amino acids [19], organic acids and sugars [13], and fatty acids [22]. Peak integration was performed using OpenChrom software [23]. Peak areas for different mass fragments corresponding to different labeling states were then used to calculate relative abundances for mass isotopomers of each analyte fragment analyzed.

### Flux Modeling

The computational platform INCA (Isotopomer Network Compartmental Analysis) [17] was used to model intracellular fluxes for *C. zofingiensis* under each condition under steady state conditions. Experimentally measured external fluxes and measured mass isotopomer distributions were used in this program along with the central metabolic network model to calculate internal flux distributions with a minimized SSR.

### Transcriptomics Experiments Strains and Culture Conditions

*Chromochloris zofingiensis* wild type strain, SAG 211-14 was grown photoheterotrophically in a modified liquid TAP medium referred to as TPNO_3_ Briefly, 7.5 mM of NH_4_Cl was replaced with 7.5 mM KNO_3_ and 17 mM acetate was replaced with 2% glucose (w/v). All glassware was pre-washed with 6 M HCl and soaked for >24 h in 6 M HCl. Flasks were rinsed with ICP-MS grade Milli-Q water (Quinn and Merchant, 1998). Ultra-pure trace metal grade chemicals and Special K micronutrient solution (PMC3101321) were used. Replete TPNO_3_ medium contained 20 µM Fe, whereas the low iron condition medium contained 2 µM Fe and the very low iron condition medium contained 0.2 µM Fe. 100 mL cultures were grown in 250 mL flasks with agitation at 180 rpm at 25 °C in an Infors HT Multitron Pro incubator, under continuous illumination of 100 µmol photons m^−2^ s^−1^ provided by warm white (3435 K) LED lights. The inoculum cultures were grown to the exponential growth phase in a replete medium and cells were washed twice in Milli-Q water before inoculation into the desired medium. Cells were counted using a Neubauer hemocytometer and all experimental cultures were inoculated to an initial density of 10^5^ cell mL^−1^. For each condition, four replicates (individual flasks referred to subsequently as biological replicates) were grown simultaneously and were sampled for RNA at the exponential growth phase (2–4 × 10^7^ cell mL^−1^).

### Nucleic Acid Extraction and Sequencing

Total RNA was extracted using a modified phenol chloroform extraction protocol. Approximately 10^8^ cells were collected by centrifugation at 3220 ×*g* for 3 min. The supernatants were discarded, and the cell pellets were resuspended in 1 mL of Tri Reagent and transferred to a 1.5 mL screwcap vial. One 4 mm glass bead and 200 mg of 0.5 mm glass beads were added to each vial prior to bead beating for 5 min using a Biospec MiniBead Beater 16. The mixtures were incubated on ice for 5 min and 200 µL of chloroform was added. The vials were vortexed for 5 s and incubated on ice for 5 min. Each sample was centrifuged for 15 min at 16,000 ×*g*, 4°C and the top aqueous layer was transferred to a new 1.5 mL tube. 500 µL of isopropanol was added to the tube with the aqueous layer and incubated overnight at −20°C. Each sample was centrifuged at 21100 ×*g*, 4 °C for 30 min and the supernatant was removed. The pellet was washed twice with fresh ice cold 80% EtOH and centrifuged at 21100 ×*g*, 4 °C for 15 min. EtOH was carefully removed, and the pellet was resuspended in 400 µL of RNase free water at room temperature for 1 hr. 50 µL of 3 M sodium acetate, pH 8 was added and the tube was inverted. 1 mL of 100% EtOH was added, and the RNA was purified using the Qiagen miRNeasy Mini Kit (Cat. No. 217004) following the manufacturer’s instructions. Purified RNA quality was assessed using the Agilent 6000 Pico Kit (Cat. No. 5067-1513) on a Bioanalyzer (Agilent, Model: 2100) and quantified using a Qubit fluorometer (Thermofisher Scientific). RNA-seq libraries were prepared and sequenced by the U.S. Department of Energy Joint Genome Institute (JGI). Stranded libraries were created from poly(A) selected transcripts using standard kits and protocols from KAPA Biosystems (Wilmington, MA) and sequenced on the Illumina NovaSeq 6000 sequencer using the NovaSeq XP V1.5 reagent kits and S4 flow cell. The SAG 211-14 genome (v.5.3) [24] with associated annotations [25] was used as a reference for the alignment of filtered reads using the splice-aware aligner STAR (v2.7.10b) with --alignIntronMax 3000 -- outSAMtype BAM SortedByCoordinate --outMultimapperOrder Random --outSAMmultNmax 1. The R package DESeq2 (v.1.44) was used for differential gene expression analysis. Genes were considered differentially expressed (DEGs) if they exhibited a log_2_-transformed fold change of >1 or <−1 and a Benjamini-Hochberg adjusted p-value of <0.01. Fragments per kilobase per million mapped reads (FPKM) values were estimated using Cuffdiff (v2.0.2).

## Results

### Culture Growth and External Flux Measurements

Cultures grown on TGP medium reach stationary phase at a lower culture density than those grown in CORE medium (Figure 2A). TGP medium had a starting iron concentration of approximately 3 µM, while CORE medium had an initial iron concentration of approximately 250 µM. By mid-exponential phase, this decreased to 0.76 µM and 63 µM in TGP and CORE media, respectively (Figure 2B). TGP cultures grown in continuous light degrade chlorophyll and they appear orange due to the production of astaxanthin (Figure 2C). Cells grown in replete CORE medium in the dark degrade their chlorophyll as well, but when grown in continuous light they retain it.

**Figure 2.**
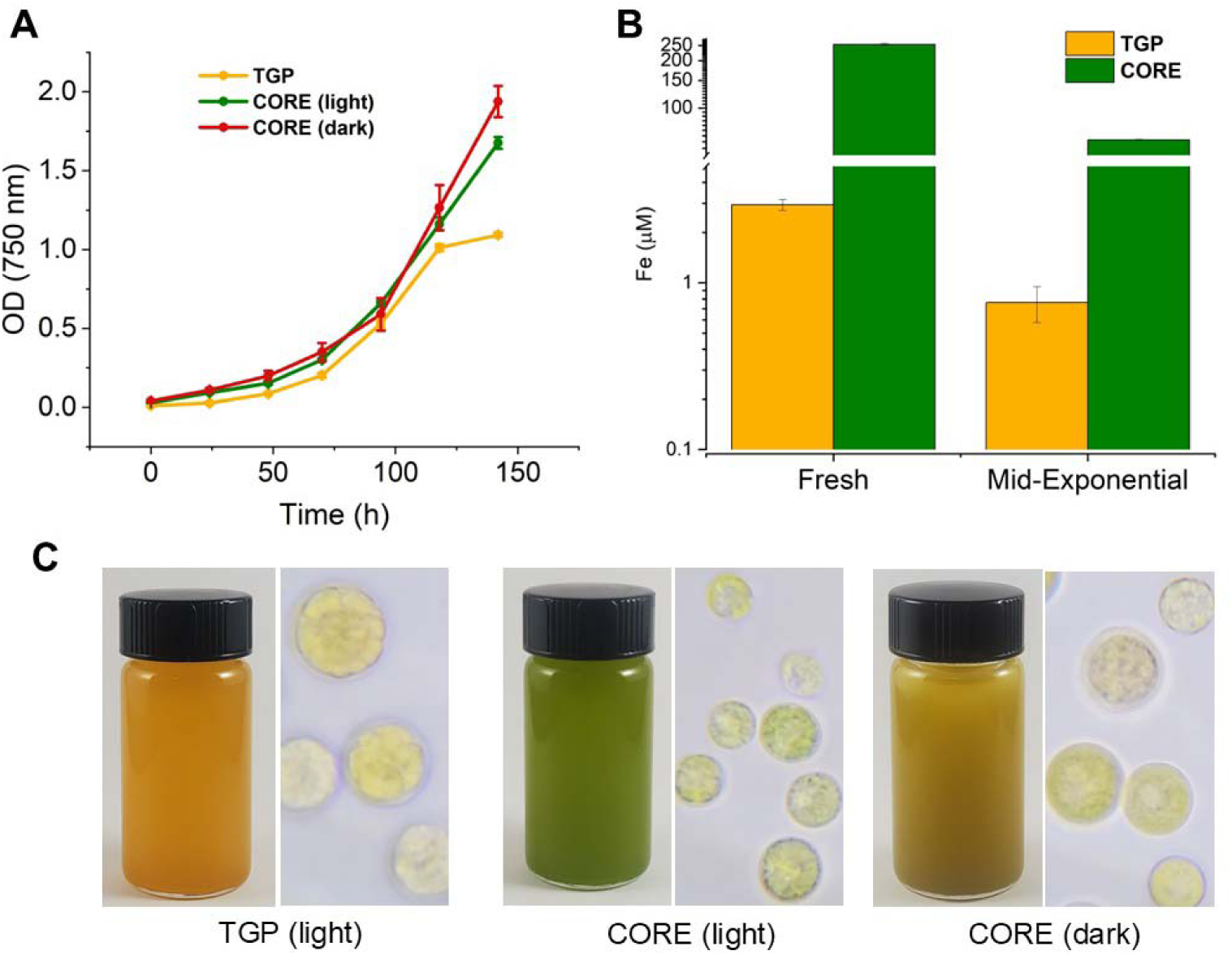
Culture growth for each condition. Panel A shows growth curves based on optical density readings for each condition (λ=750nm). Error bars represent the standard deviation of n=3 biological replicates. Panel B shows initial and mid-exponential iron concentrations for TGP and CORE media in continuous light. Iron measurements were carried out using the Cedex bioanalyzer iron assay. Error bars represent the standard deviation of n=3 biological replicates. Panel C shows photographs of *C. zofingiensis* cultures grown in TGP media in continuous light, CORE media in continuous light, and CORE media in continuous dark conditions.

Samples of filtered media collected during growth were analyzed to measure glucose, lactate, and succinate concentrations over time. These data were then used to calculate uptake and excretion fluxes of glucose, lactate, and succinate in TGP media in continuous light, and CORE media in continuous light and in dark growth conditions. Cultures grown on TGP media exhibited the highest glucose uptake flux and were the only cultures where fermentation product secretion was observed (Figure *3*). CORE media cultures grown in continuous light had the lowest glucose uptake flux, as expected for mixotrophic growth (Figure *3*).

**Figure 3.**
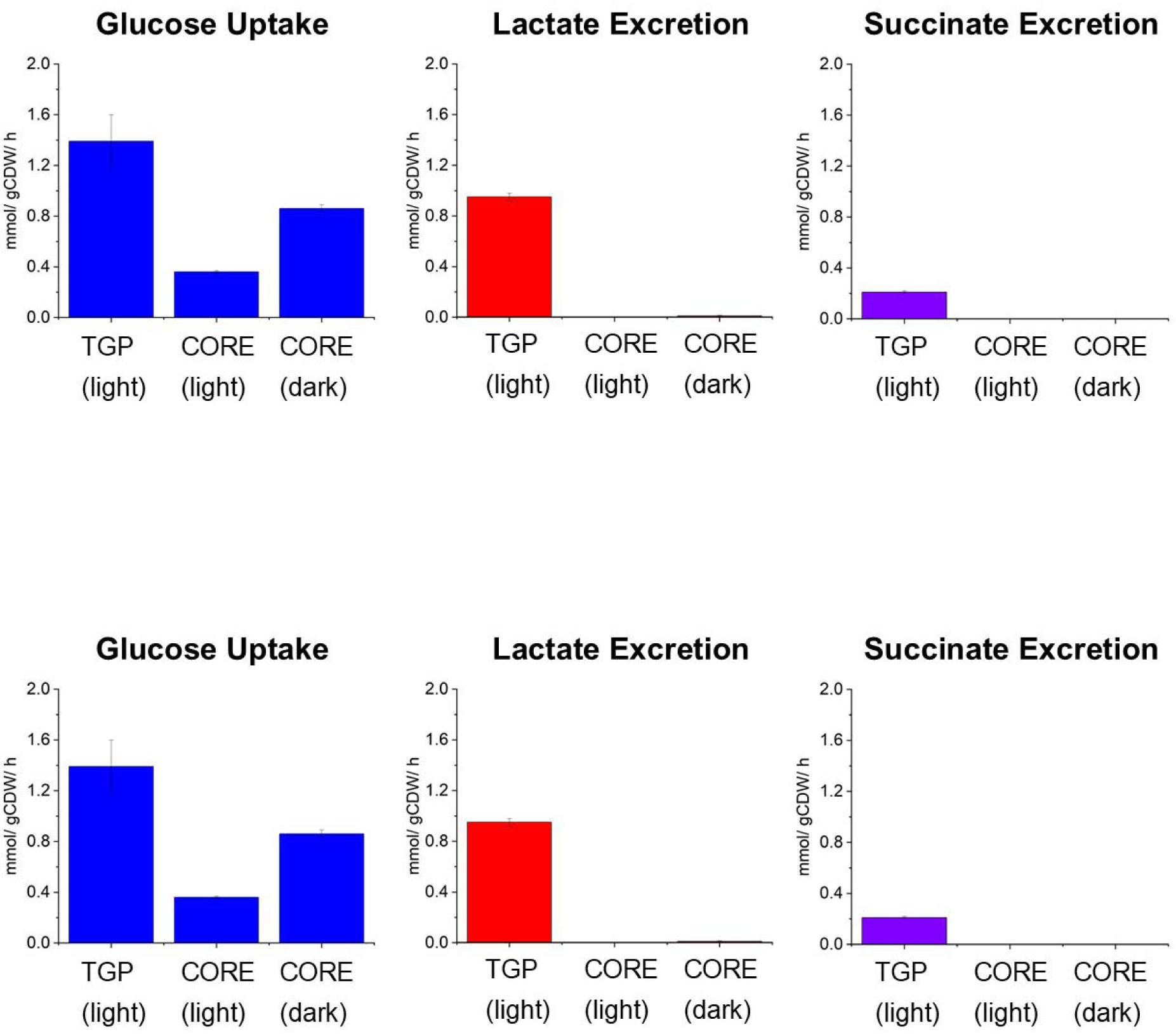
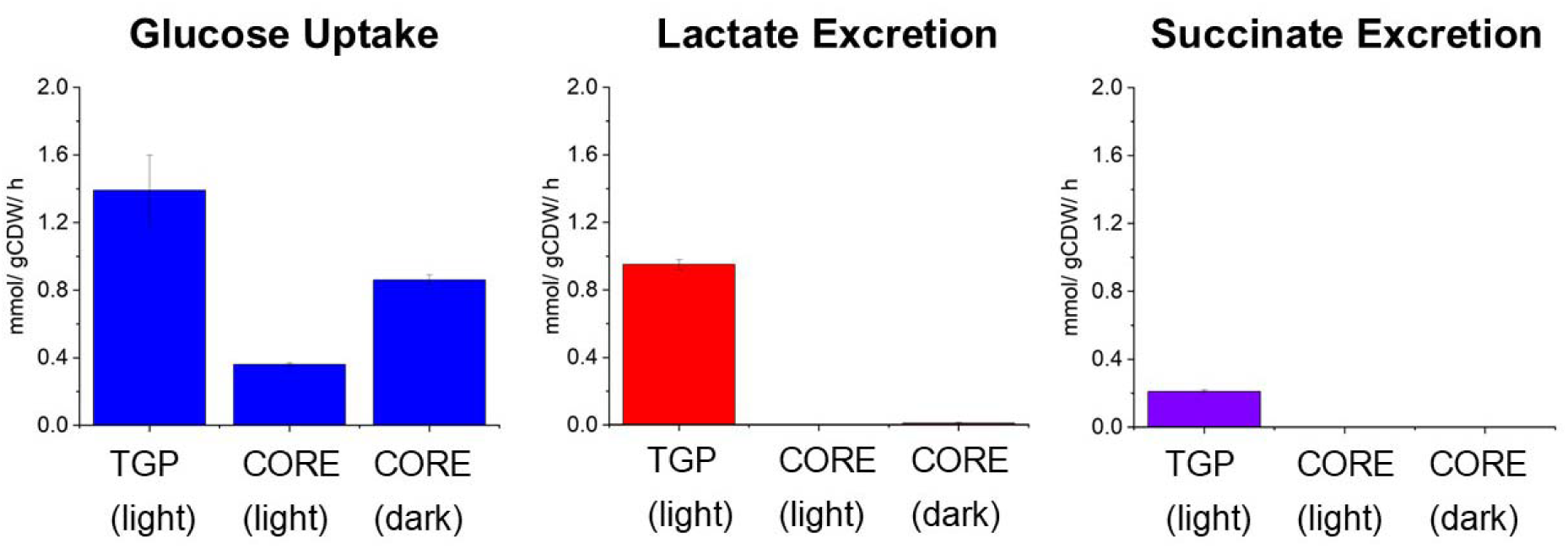
External fluxes used for metabolic flux analysis. Glucose uptake and fermentation product secretion rates resulting from analysis of spent media samples collected during growth experiments for each condition in this experiment. Error bars represent standard deviation of n=3 biological replicates. The full data set used to calculate exchange fluxes is presented in supplemental figure S1.

### Central Metabolic Network Model

The genome-scale metabolic model (GSM) of *C. zofingiensis*, *i*Czof1915 [10], was used to construct a smaller model of central carbon metabolism in three cellular compartments: cytosol, chloroplast, and mitochondria. Reactions comprising the major central metabolic pathways glycolysis, TCA cycle, pentose-phosphate (PPP) and Calvin-Benson-Bassham (CBB) cycle, as well as fermentation, and amino acid, carbohydrate, and fatty acid biosynthesis were included (see Figure 4). The addition of certain secondary metabolites into the network was necessary in this compartmentalized model, as compartment specific labeling information can be obtained for carbohydrates, amino acids, and fatty acids [26]. Amino acid biosynthesis pathways were included only for amino acids whose mass fragments have been reported to be reliable for metabolic flux analysis [19]. The amino acids asparagine and glutamine are deaminated during the sample derivatization process before analysis on GC-MS [19], so these amino acids are grouped in with aspartate and glutamate, respectively. The final list of amino acids in this metabolic network included alanine, aspartate, glutamate, glycine, isoleucine, leucine, methionine, phenylalanine, serine, threonine, tyrosine, and valine. Carbohydrate biosynthesis pathways included were the production of starch and cell wall sugars galactose and arabinose. Because the cells are actively growing during the time course of these labeling experiments, the network model had to account for the formation of biomass. This was done by providing a sink reaction for each biomass component, such as amino acids and fatty acids. In order to account for the production of other biomass components not explicitly formed in the network, a sink reaction was added for each of the 12 central metabolism components of biomass [27]. After the central network of reactions had been pulled together, atom transitions for each reaction were assigned based on associated KEGG enzymes for each reaction [28]. The metabolic network was then condensed to collapse linear reaction pathways, easing computational load for isotopic labeling simulations. The final condensed network model consisted of 96 reactions, including 62 metabolic reactions, 12 transport reactions, and 22 reactions to account for metabolite sinks for biomass formation. This reaction network is shown in Figure 4. A full list of all metabolite abbreviations can be found in supplementary table S1. A list of all reactions and atom transitions included in the model can be found in table S3. In Figure 4, analytes whose mass isotopomer distributions were measured are highlighted in blue. Data on mass isotopomer distributions (MID) measured via GC-MS for amino acids, organic acids, fatty acids, and sugars are displayed in supplementary figures S2-S5.

**Figure 4.**
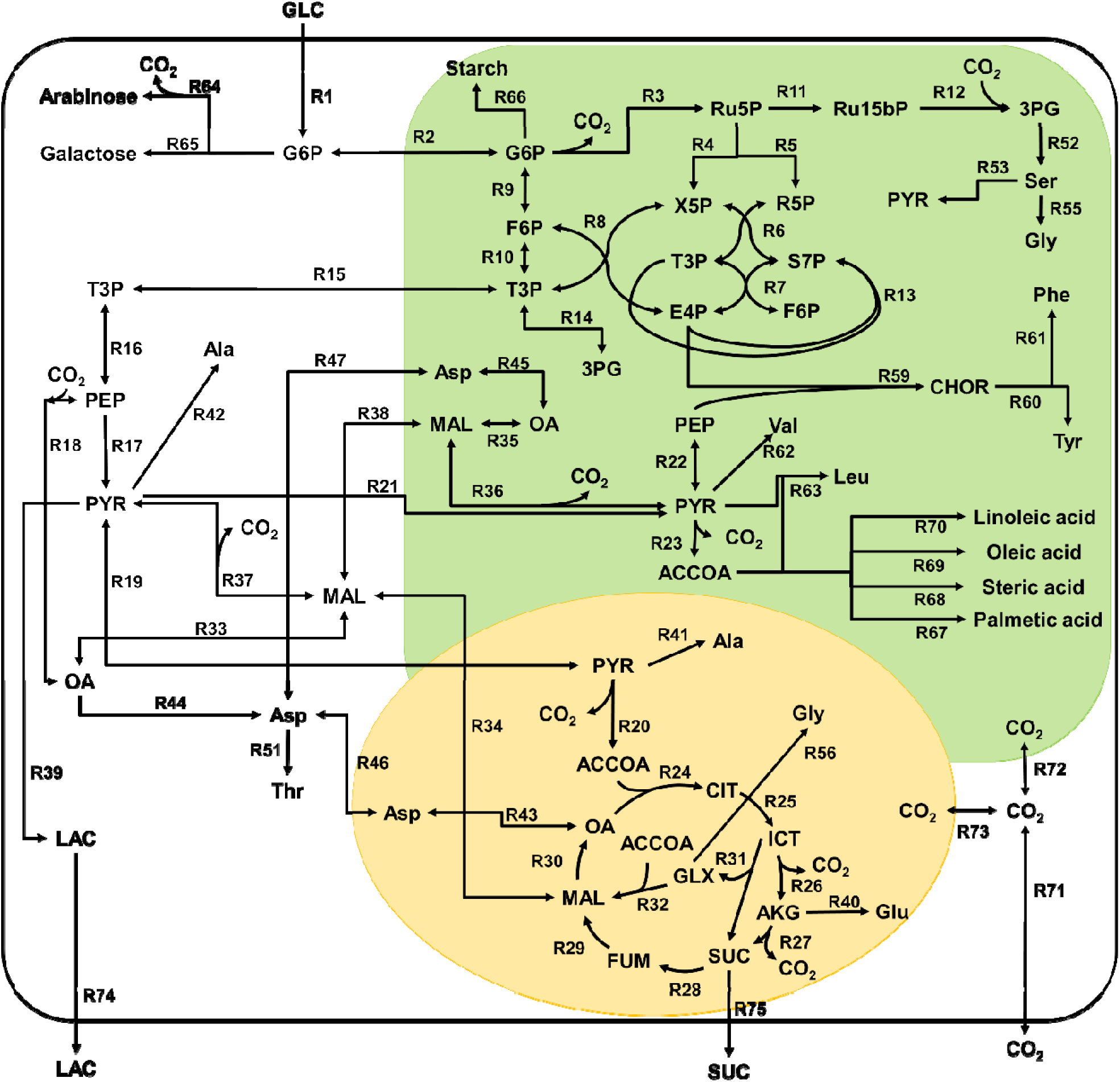
Central metabolic network model of *C. zofingiensis.* This reaction network model consists of three compartments (cytosol in white, chloroplast in green, and mitochondria in yellow). A full list of metabolite abbreviations is found in Supplemental Table S1. A full list of mass isotopomer fragments measured is found i Supplemental Table S2. A full list of reactions and atom transitions included in the model is found in Supplemental Table S3.

### Intracellular Flux Distributions

Calculated intracellular flux distributions for TGP culture grown in light, CORE cultures grown in light, and CORE cultures grown in dark are shown in **Error! Reference source not found.**, **Error! Reference source not found.**, and **Error! Reference source not found.** respectively. For ease of comparison between each case, fluxes have been normalized to a glucose uptake of 100 mmol/grams cell dry weight (gCDW)/h. In TGP cultures (**Error! Reference source not found.**), after uptake, glucose is directed into the plastid and is directed into either glycolysis or the pentose phosphate pathway. No flux occurs through carbon fixation pathways, consistent with previous observation of low Fv/Fm (0.01) measured for this phenotype of *C. zofingiensis* [2]. Carbon is exported from the plastid in the form of triose phosphate sugars. The latter half of glycolysis proceeds in the cytosol until PEP, at which point approximately 40% of the carbon flux is diverted through anapleurotic reactions via PEP carboxylase rather being processed to pyruvate. The carbon flux that does flow through pyruvate is then split with 60% of the flux directed to lactate formation, and 40% directed to the mitochondria to enter the TCA cycle. In TGP cultures, the full TCA cycle is not used. The cell relies heavily on the glyoxylate shunt and there is no predicted flux through the reaction catalyzed by isocitrate dehydrogenase.

**Figure 5.**
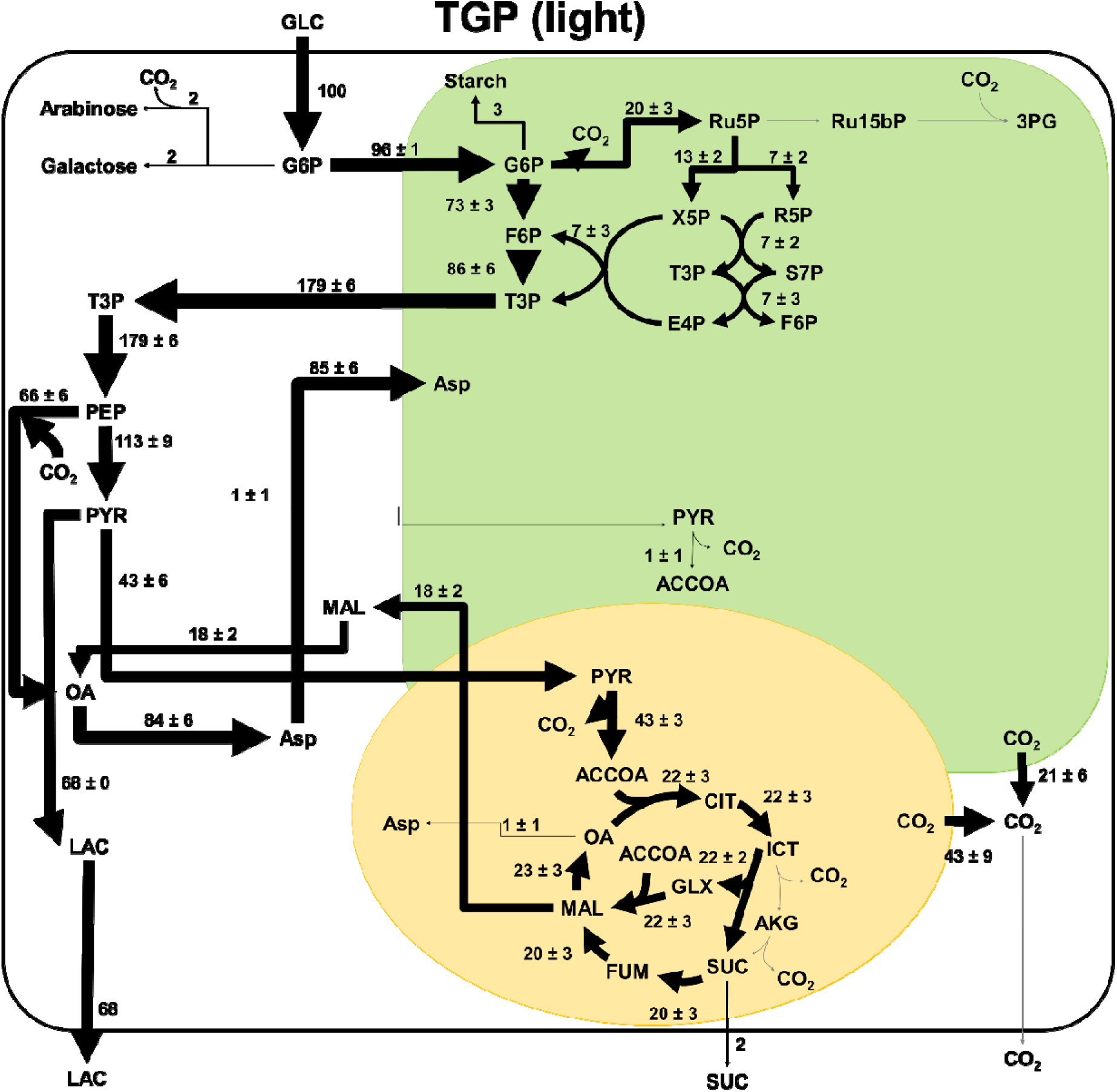
Intracellular fluxes for TGP cultures. Fluxes have been normalized to a glucose uptake flux of 10 mmol/gCDW/h. Fluxes with negligible values are shown in grey. This reaction network model consists of three compartments (cytosol in white, chloroplast in green, and mitochondria in yellow). Arrow thickness is scaled to the magnitude of flux through each reaction. A full list of metabolite abbreviations is found in Supplemental Table S1.

In CORE in light cultures (**Error! Reference source not found.**), after uptake, glucose is directed into the plastid and progresses through the initial steps of glycolysis. Very little flux i observed through the oxidative phase of the pentose phosphate pathway, while in comparison the reductive pentose phosphate pathway reactions and the Calvin-Benson-Bassham cycle are more active. This is the only case studied where *C. zofingiensis* maintains photosynthetic activity while growing on glucose, and is the only case with non-negligible flux through the Calvin Benson Bassham Cycle. Carbon is exported from the plastid as triose phosphate sugars, which then progress through glycolysis to pyruvate. In these cultures, negligible flux was observed through PEP carboxylase, with all of the carbon flowing through pyruvate kinase into pyruvate. From here, around 15% of the carbon flux re-enters the plastid as pyruvate at which point this pyruvate participates in a cycle where malate is formed in the plastid consuming NADH, then this malate is exported to the cytosol and converted back to pyruvate, producing NADH before being transported back to the plastid. This cycle functions mainly to move reducing equivalents from the plastid to the cytosol. The remaining carbon that is not participating in this cycle is sent to the TCA cycle. In these cultures, carbon flux was found through both the glyoxylate shunt and the full TCA cycle.

**Figure 6.**
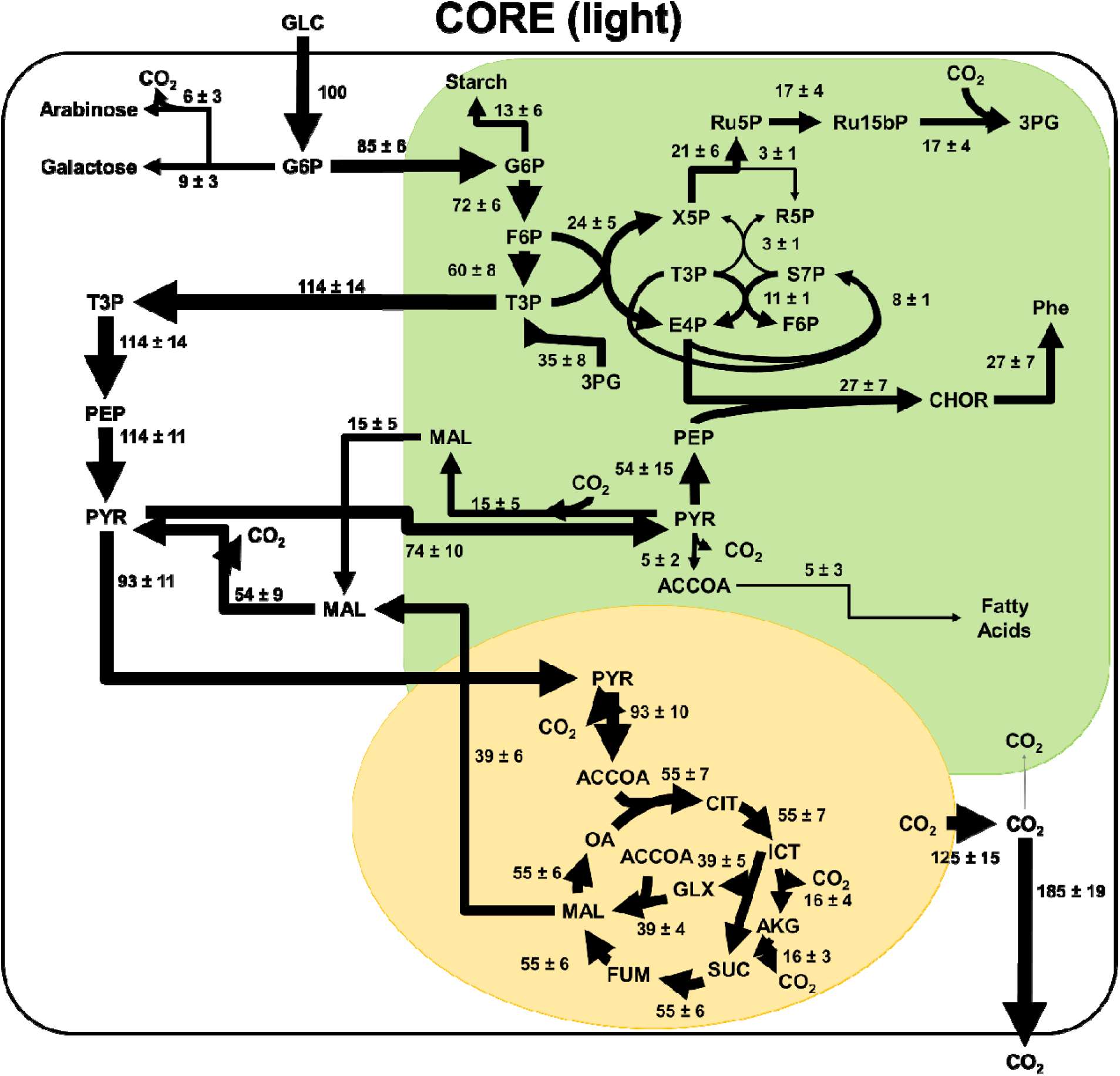
Intracellular fluxes for CORE cultures grown in light. Fluxes have been normalized to a glucose uptake flux of 100 mmol/gCDW/h. Fluxes with negligible values are shown in grey. This reaction network model consists of three compartments (cytosol in white, chloroplast in green, and mitochondria in yellow). Arrow thickness is scaled to the magnitude of flux through each reaction. A full list of metabolite abbreviations is found in Supplemental Table S1.

In CORE dark cultures (**Error! Reference source not found.**), after uptake, glucose is sent to the plastid, where carbon flux is split between glycolysis and the pentose phosphate pathway. Carbon is exported from the plastid in the form of triose phosphate sugars, where it continues through the lower steps of glycolysis until phosphoenolpyruvate. At this point, a majority of the flux is directed through PEP carboxylase forming oxaloacetate, which is then converted to malate. A large portion of this malate is then converted to pyruvate through malic enzyme. The carbon flux from pyruvate is split between the mitochondria where it enters the TCA cycle, and the plastid where the carbon flux is either directed to the formation of biomass components or is converted back to malate.

**Figure 7.**
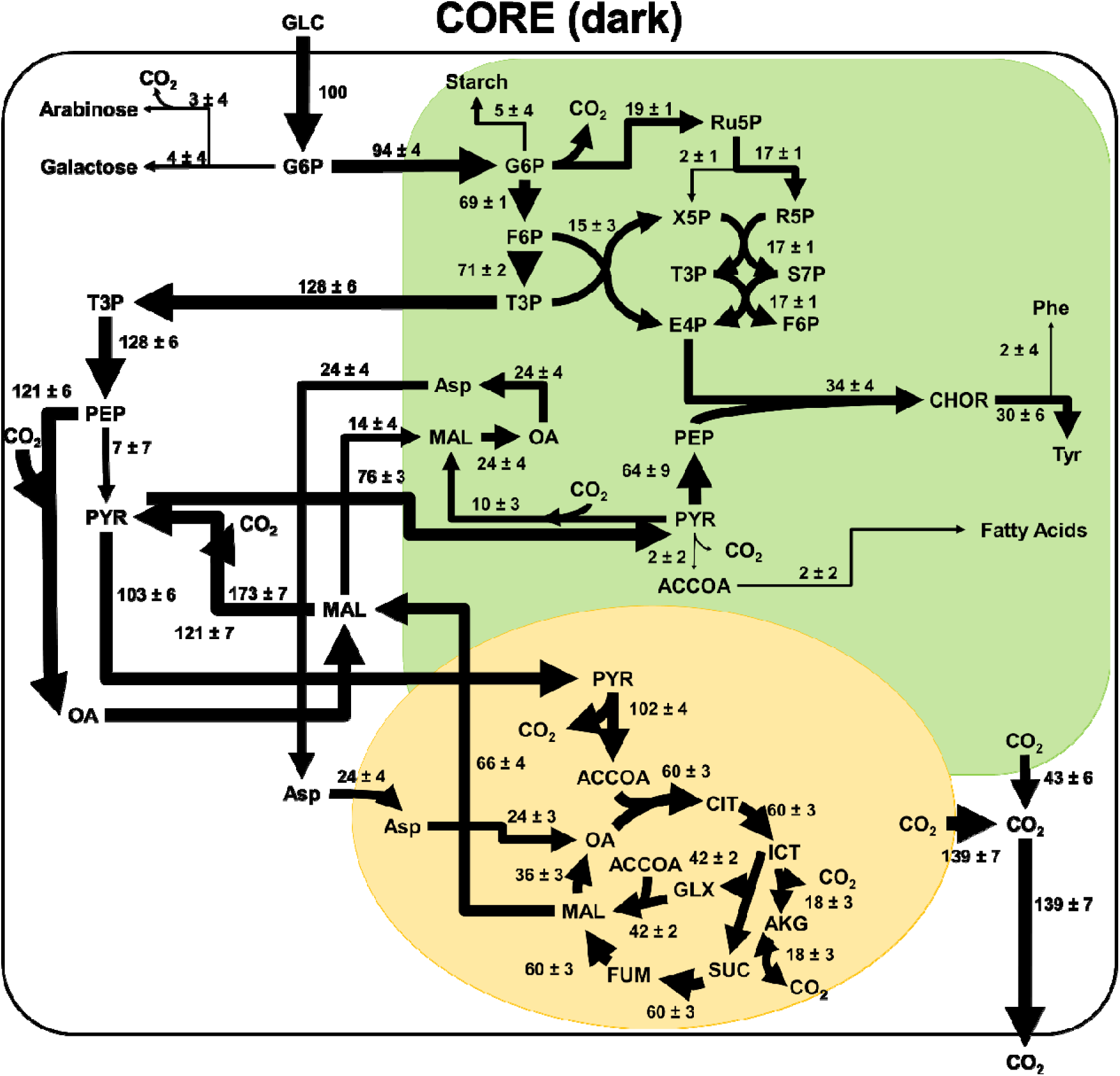
Intracellular fluxes for CORE cultures grown in the dark. Fluxes have been normalized to a glucose uptake flux of 100 mmol/gCDW/h. Fluxes with negligible values are shown in grey. This reaction network model consists of three compartments (cytosol in white, chloroplast in green, and mitochondria in yellow). Arrow thickness is scaled to the magnitude of flux through each reaction. A full list of metabolite abbreviations is found in Supplemental Table S1.

### Transcriptomics Data

Transcript abundance for low and very low iron conditions was compared to the iron replete condition to calculate log_2_ fold changes in gene expression. The full set of transcriptomics data can be found in supplemental file S1. The change in expression for genes associated with enzymes in the TCA cycle localized in the mitochondria, and the glyoxylate shunt localized in the peroxisome are found in Figure 8. Transcripts for the gene associated with the isocitrate lyase enzyme were identified in low abundance and had little change in expression for the low and very low iron conditions. However, the gene Cz14g25060 which is annotated as encoding for malate synthase was found to have a higher expression in the low and very low iron growth conditions (Figure 8). The genes Cz07g14020 and Cz15g13230, which are associated with fumarase, also had an increase in expression levels in low iron conditions (Figure 8). Other than these examples, the majority of genes associated with TCA cycle enzymes had lower expression levels in low iron growth conditions.

**Figure 8.**
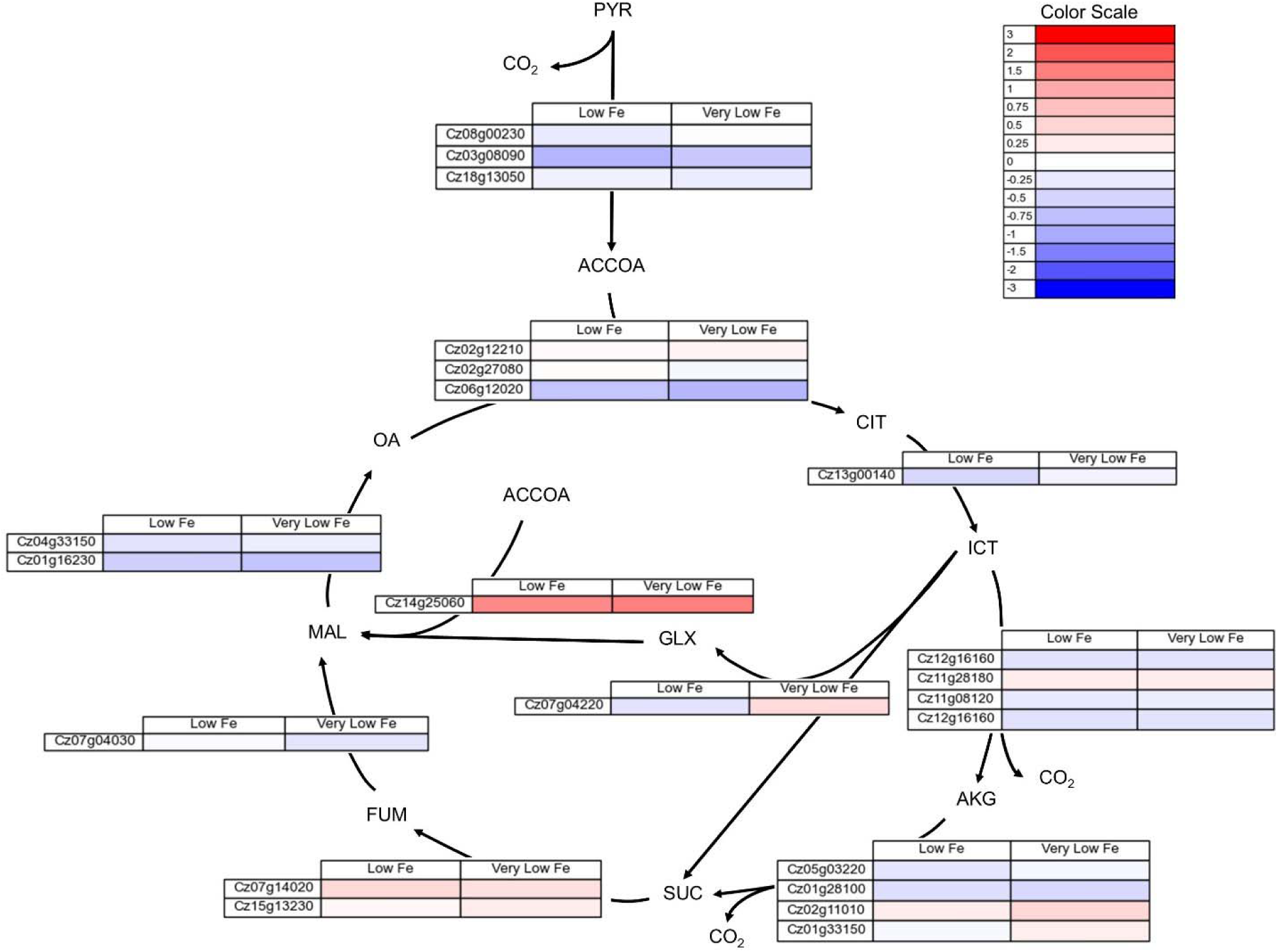
Log_2_ fold change in gene expression for genes associated with TCA cycle enzymes localized to the mitochondria and glyoxylate shunt enzymes localized to the peroxisome for low (2 µM Fe) and very low (0.2 µM Fe) iron as compared to the replete (20 µM Fe) condition. The full set of FPKMs and log_2_ fold change data is available in supplemental table S4.

The changes in expression for genes associated with Calvin Benson-Bassham (CBB) cycle enzymes are found in Figure 9. The majority of genes associated with CBB cycle enzymes have lower expression in low and very low iron conditions, particularly the genes associated with RuBisCo.

**Figure 9.**
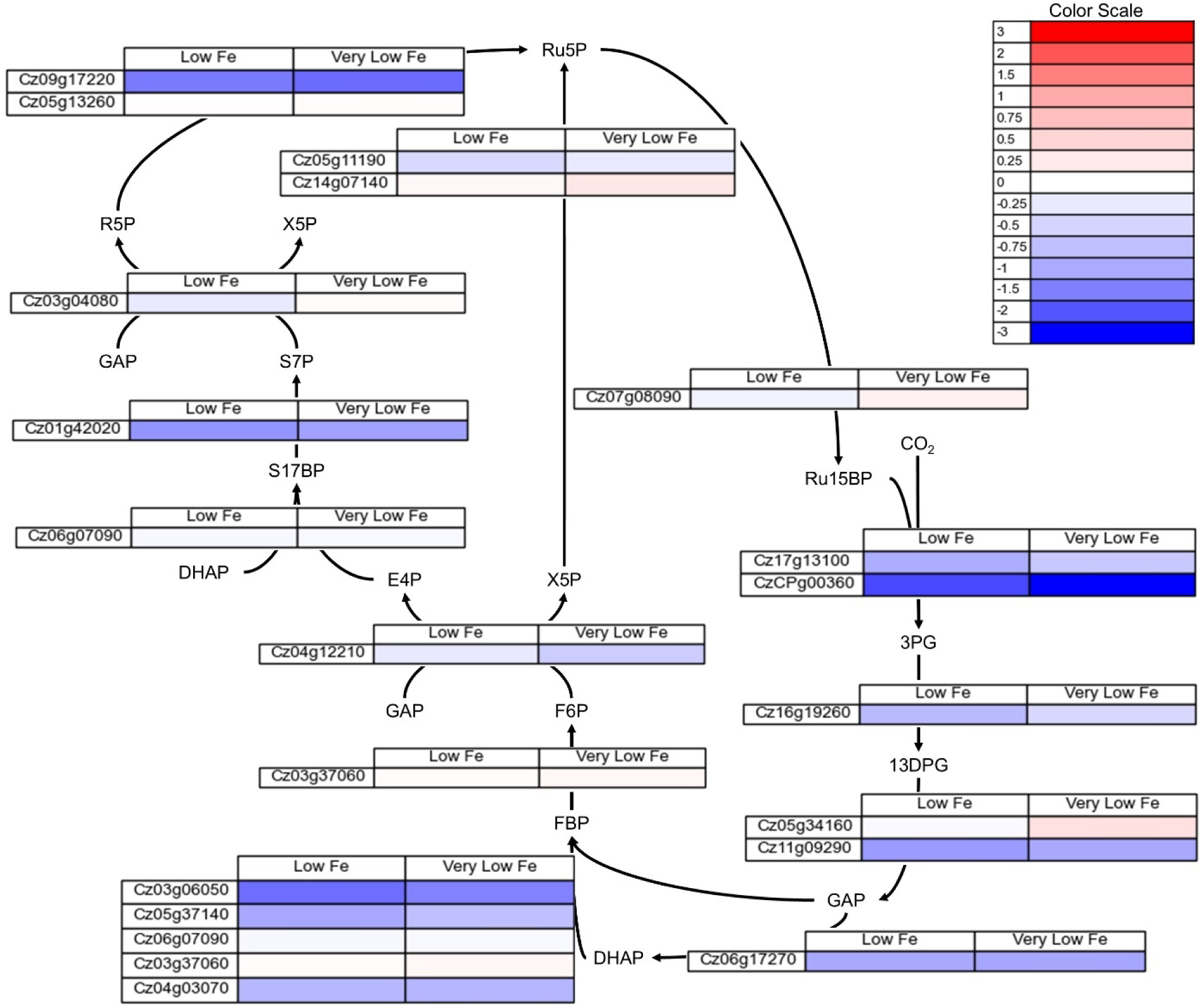
Log_2_ fold change in gene expression for genes associated with CBB cycle enzymes localized to the chloroplast for low (2 µM Fe) and very low (0.2 µM Fe) iron as compared to the iron replete condition. The full set of FPKMs and log_2_ fold change data is available in supplemental table S5.

In addition to genes associated with metabolic enzymes, genes associated with photosynthetic and respiratory electron transport chains were analyzed.

Figure **10** shows the log_2_ fold change in genes associated with the respiratory electron transport chain and ATP synthase in the mitochondria for low and very low iron conditions. Figure 11 shows the log_2_ fold change in genes associated with the photosynthetic electron transport chain and chloroplastic ATP synthase. There is a decreased expression of genes associated with both the respiratory and photosynthetic electron transport chains in low and very low iron conditions. The genes for both the mitochondrial and the chloroplast ATP synthase also have lower expression in low and very low iron conditions, indicating that growth in low iron media impacts the ability of *C. zofingiensis* to generate ATP.

**Figure 10.**
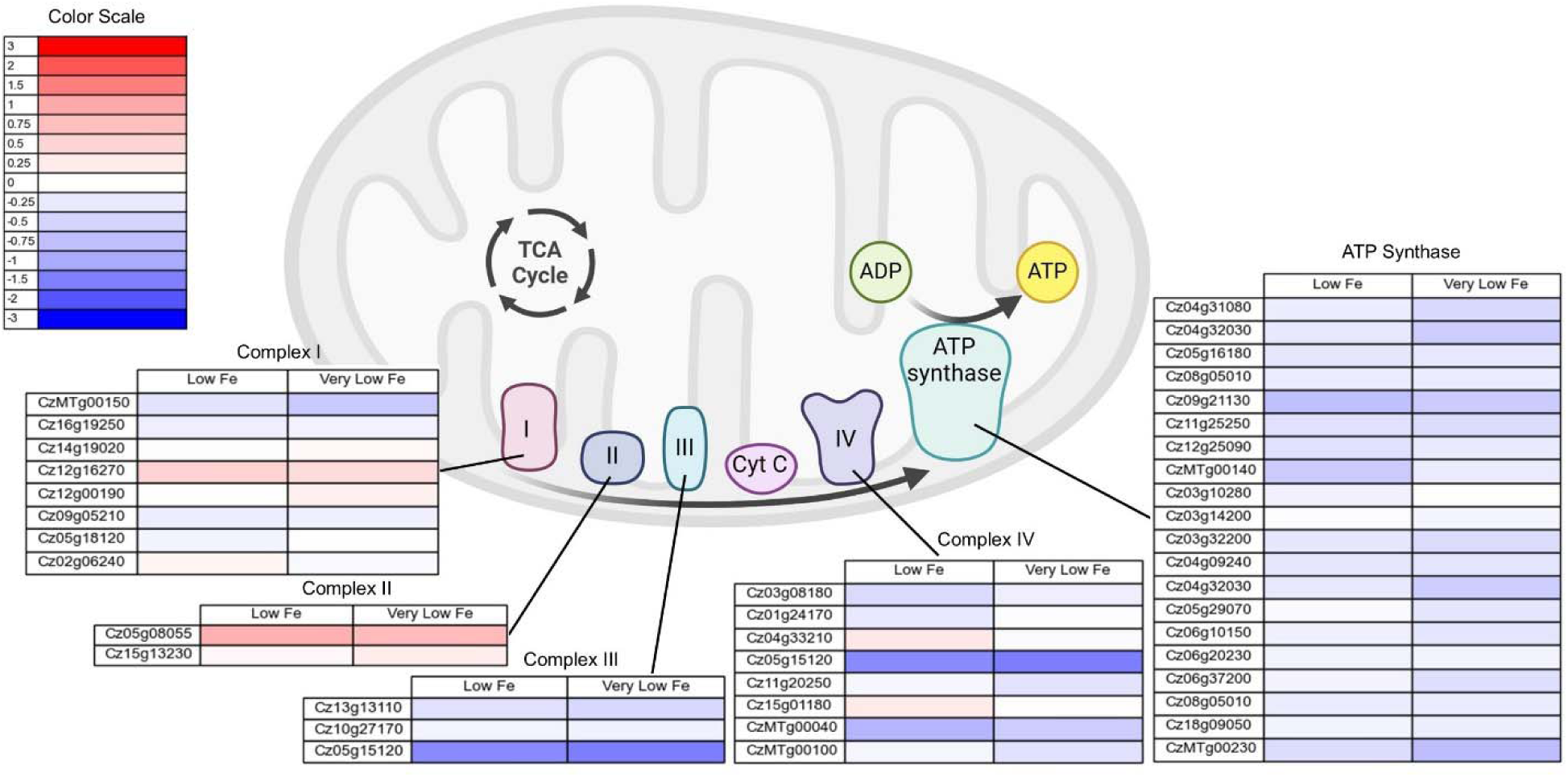
Log_2_ fold change in expression of genes associated with the mitochondrial respiratory electron transport chain and ATP synthase for low (2 µM Fe) and very low (0.2 µM Fe) iron conditions. Figure was made using Biorender. The full set of FPKMs and log_2_ fold change data is available in supplemental table S6.

**Figure 11.**
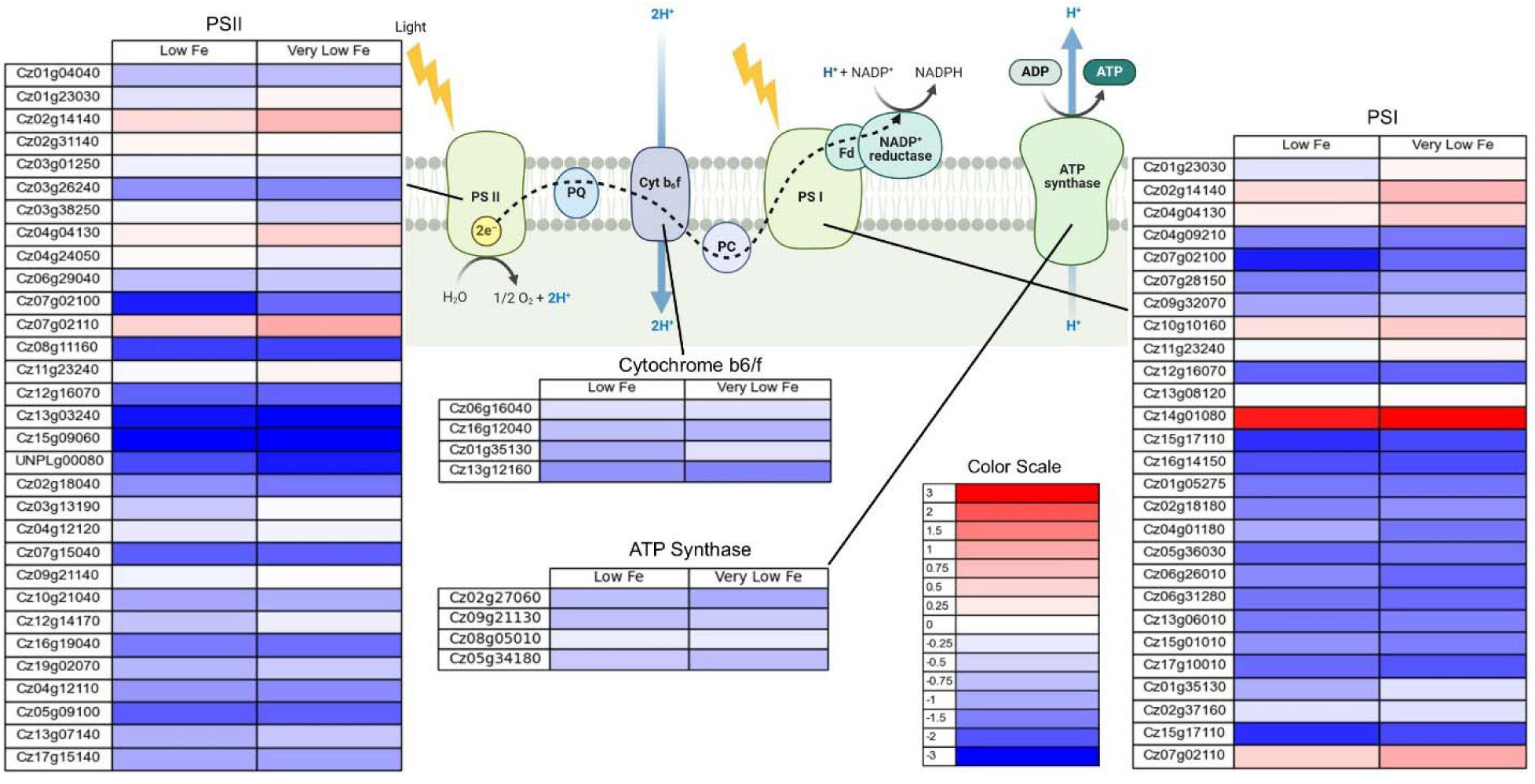
Log_2_ fold change in expression of genes associated with the photosynthetic electron transport chain and chloroplastic ATP synthase for low (2 µM Fe) and very low (0.2 µM Fe) iron conditions. Figure was made using Biorender. The full set of FPKMs and log_2_ fold change data is available in supplemental table S7.

## Discussion

### Culture Growth

TAP medium is considered the standard for cultivation of the model green alga *C. reinhardtii*, and as such it is frequently used in the cultivation of other green algae species. However, the nutrients present in TAP are not necessarily optimal for all green algae. As a soil alga [29], *C. zofingiensis* appears to have very different nutrient requirements for optimal growth than the planktonic *C. reinhardtii*. The comparison of *C. zofingiensis* growth in TGP versus nutrient-optimized CORE medium highlight the fact that large metabolic changes can result from the use of a culture medium with sub-optimal nutrient concentrations. The largest differences between TGP and CORE media are the amount and type of nitrogen provided (NH_4_ in TGP vs. NO_3_ in CORE), and the levels of iron and magnesium. Iron, in particular, was two orders of magnitude higher in CORE media than in TGP media (Figure 2B). The expected iron concentration of TGP medium based on media recipes is around 18 µM, however, when iron was measured retrospectively in fresh TGP medium it was found to contain only around 3 µM Fe. This is likely a consequence of the use of Hutner’s trace element mixture in the preparation of TGP medium, as iron precipitation occurs in the preparation of this mixture leading to lower than expected iron concentrations and variability in metal concentrations between batches [30]. Magnesium was also an order of magnitude lower in TGP medium than in CORE, and Mg^2+^ plays a vital role as a cofactor for a number of enzymes. These differences in nutrient availability translate into very different metabolic phenotypes. Table 1 below examines the differences in growth rates and biomass productivities on glucose for each cultivation condition examined in this work. In CORE media, higher growth rates and higher biomass productivities are seen for cultures grown in continuous light than those grown in the dark, which is expected due to the additional energy provided to the cell by light. Growth in TGP media in the light results in the highest growth rate and the lowest biomass productivity on glucose. TGP media growth was also the only condition where fermentation product secretion was observed. If this fermentation was due to a lack in oxygen availability, it would be expected in the CORE dark cultures as well, as these cultures were grown heterotrophically in the same baffled flasks as the TGP cultures. Because fermentation was not observed in any CORE cultures, it is evident that this fermentation of glucose occurs due to the differences in media composition between the two media types. Additionally, previous work has shown that when *C. zofingiensis* is grown heterotrophically on TGP media in a steady-state continuous flow bioreactor, lactate excretion still occurs despite the oxygen levels within the bioreactor remaining above the threshold for anoxic conditions [10].

**Table 1.** Summary of growth rates and biomass productivities for each growth condition. Growth rates were evaluated using cultivation data shown in Figure 2.

| Growth Condition | Growth Rate (h<sup>-1</sup>) | Biomass Productivity |
| --- | --- | --- |
|  |  | (g CDW produced/ g glucose consumed) |
| TGP in light | 0.041 ± 0.001 | 2.6 e-2 ± 1e-3 |
| CORE in light | 0.031 ± 0.001 | 9.5 e-1 ± 8e-5 |
| CORE in dark | 0.028 ± 0.001 | 6.7 e-2 ± 4e-4 |

### Flux Distributions

In this network model, there are two major pathways that carbon flux could be directed through phosphoenolpyruvate. Pyruvate kinase converts phosphoenolpyruvate to pyruvate in the last step of glycolysis, and PEP carboxylase converts phosphoenolpyruvate and CO_2_ to oxaloacetate in a reaction that typically serves an anapleurotic function replenishing TCA intermediates. A higher flux through PEP carboxylase was observed for both TGP grown in the light and CORE dark cultures, while in CORE light cultures the flux through this enzyme was negligible, with flux entirely directed through pyruvate kinase. In CORE dark cultures, there was negligible flux through pyruvate kinase and the cell relies entirely on PEP carboxylase, using this enzyme as well as malate dehydrogenase and malic enzyme to eventually produce pyruvate. In plants, it has been shown that pyruvate kinase is deactivated in dark conditions [31] [32]. This light-based inactivation is a likely explanation for why this enzyme is inactive in CORE cultures grown in the dark. This route of carbon into the TCA cycle through PEP carboxylase and malic enzyme to generate pyruvate rather than pyruvate kinase has been observed in other modeling studies in heterotrophic algae [34-38]. In TGP cultures, carbon flux is split between PEP carboxylase and pyruvate kinase, with approximately 40% of the total flux going toward oxaloacetate synthesis. This diversion of flux away from pyruvate kinase comes at the cost of ATP formation. Pyruvate kinase catalyzes the last step in glycolysis, and controls the flow of carbon into the TCA cycle, which is used for both respiratory metabolism and to provide metabolic intermediates used in biomass synthesis. By controlling the flow of carbon into the TCA cycle, pyruvate kinase controls the balance between these two metabolic paths [33]. When comparing flux between TGP and CORE light cultures, TGP appears to have a down-regulation of carbon entering the TCA as evidenced by a diversion of flux through PEP carboxylase.

Another difference in flux distributions between culture conditions is seen in the pentose phosphate pathway. TGP cultures and CORE cultures grown in the dark have a higher flux through the oxidative steps of the pentose phosphate pathway, whereas CORE cultures grown in the light have a negligible flux through these reactions. The oxidative pentose phosphate pathway is important to heterotrophic algae growth as it provides NADPH production in the absence of photosynthesis. When comparing pentose phosphate pathway fluxes between both heterotrophic growth cases, CORE cultures grown in the dark had a higher flux through the whole of the pentose phosphate pathway. In *C. reinhardtii*, it has been found that Mg^2+^ is required for optimal function of the transketolase enzyme [40], responsible for catalyzing reactions in the non-oxidative phase of the pentose phosphate pathway. It is possible that the lower Mg^2+^ concentration in TGP media contributes to the difference in pentose phosphate pathway activity between TGP and CORE cultures.

### Photosynthesis Activity

Metabolic flux analysis results show that CORE media cultures grown in continuous light have flux through carbon fixation pathways, while TGP cultures grown in the same conditions do not. Previous work has shown the importance of iron for photosynthetic activity in *C. zofingiensis* [3], so it is likely that the low iron content of TGP media is responsible for the loss of photosynthetic activity in TGP cultures.

This finding is reinforced when examining the expression levels for genes associated with enzymes involved in the CBB cycle. In low and very low iron conditions, the expression levels of most genes encoding for enzymes in the CBB cycle are decreased, particularly RuBisCo (Figure 9). Low iron conditions also led to a decrease in the expression of genes associated with the photosynthetic electron transport chain and the chloroplastic ATP synthase (Figure 11). Photosystems I and II were particularly impacted. One gene, Cz14g01080, had a strong increase in expression in the low iron condition. This gene is annotated as thylakoid iron deficiency induced 1 (TIDI1), an alternate light harvesting complex protein that is up regulated in response to low Fe [41].

Other work has shown that the shutdown of photosynthesis that occurs in *C. zofingiensis* cultures during heterotrophic growth doesn’t occur when cultures are supplemented with sufficient iron [3]. It’s well known that photosynthesis is heavily dependent on iron. Iron is required in both PSI and PSII as well as in components of the photosynthetic electron transport chain [42]. An estimated 21 atoms of Fe are required for one photosynthetic electron transport chain unit [4]. Work in other photosynthetic organisms has shown that a lack of sufficient iron can lead to the downregulation of photosynthesis. In *Arabidopsis thaliana*, iron deficiency leads to a down regulation of genes associated with photosynthesis [7]. *Scenedesmus* grown in low iron conditions exhibits lower CO_2_ fixation rates and lower photosynthetic activity [5]. This has also been observed in cyanobacteria, where a moderate iron deficiency triggers a preference to flavodoxin over ferredoxin to lower iron demand, but in very low iron conditions the activity of the photosynthetic electron transport chain suffers [4]. *C. zofingiensis* also has flavodoxin, encoded by gene Cz14g01090, which is highly upregulated in low and very low iron conditions (See supplemental data file 1). While flavodoxin aids in acclimation to low iron stress, it is ultimately a less efficient electron carrier than ferredoxin [43]. In *Chlamydomonas reinhardtii*, when cells are grown with an organic carbon source on low iron, the organism prioritizes the maintenance of respiration enzymes over keeping photosynthesis active [6]. A proteomic analysis revealed that in low iron mixotrophic growth conditions, *C. zofingiensis* cultures prioritize maintaining mitochondrial Fe-dependent proteins over plastid Fe-dependent proteins [3]. In this work, other nutrients were surveyed for their effect on photosynthesis in *C. zofingiensis*, including nitrogen, phosphorus, magnesium, sulfur, manganese, copper, and zinc, but only the addition of iron was found to rescue glucose mediated photosynthesis repression [3]. This suggests that while other nutrients are required, iron is the most critical for photosynthetic activity in *C. zofingiensis*. However, it should be noted that RuBisCo requires Mg^2+^ for activation [44], so it is possible that under sufficiently low Mg^2+^ conditions, the activity of this enzyme will be impacted.

### Respiration

Iron plays an important role in mitochondrial electron transport and TCA cycle activity. The requirement for Fe-S clusters in mitochondrial electron transport chains is highly conserved across eukaryotic cells [45], and several TCA cycle enzymes have a requirement for iron. The production of one respiratory electron transport chain unit requires approximately 30 atoms of iron [46].

In both light and dark growth conditions, nutrient replete CORE cultures utilized higher flux through the TCA cycle than was observed for TGP media cultures. TGP cultures had the lowest flux through the TCA cycle, with a large flux of carbon directed instead to lactate production. TGP cultures also showed flux through an incomplete TCA cycle, utilizing the glyoxylate shunt to bypass isocitrate dehydrogenase and the alpha-ketoglutarate dehydrogenase complex. Both light and dark cultures grown in CORE media had flux through the full TCA cycle, allowing for more NADPH production in these cultures.

Most genes encoding for TCA cycle enzymes exhibit decreased expression in low iron conditions, with the exception of the gene encoding succinate dehydrogenase (Figure 8), which had a slightly higher expression in low and very low iron conditions. The gene encoding glyoxylate shunt enzyme malate synthase (Cz14g25060) also had an increase in expression in low iron growth (Figure 8). Transcripts for the gene encoding isocitrate lyase were present at less than 1 FPKM. A transcriptomics analysis of two species of heterotrophic marine bacteria in low iron conditions displayed similar results, with increased expression of genes encoding glyoxylate shunt related enzymes, while genes encoding aconitase and genes associated with oxidative phosphorylation had decreased expression [47]. In this study, expression of genes associated with succinate dehydrogenase was unchanged but interestingly genes associated with fumarase were found to have a higher expression in iron limited conditions despite its requirement for iron cofactors [47]. The TCA cycle enzyme aconitase is particularly iron intensive, and requires a 4Fe-4S cluster for activity [48]. In extracts from mammalian cells, it has been shown that low iron conditions leads to lower activities of citrate synthase, aconitase, isocitrate dehydrogenase, and succinate dehydrogenase [49].

In our analysis, the genes associated with respiratory electron transport chain complex I had limited changes in expression levels in low iron conditions, and genes associated with complex II had a slight increase in expression (Figure **10**). Genes associated with complexes III and IV had lower expression levels in low iron conditions, as did the genes associated with mitochondrial ATP synthase (Figure **10**). In *Arabidopsi*s roots, iron limitation leads to a decrease in specific activity of every complex in the respiratory electron transport chain, and the genetic response to iron or sulfur limitation presents similarly to the genetic response seen in plants with mitochondrial respiration impairment [8]. While there are examples of iron limitation in algae in the literature, the vast majority of these studies focus on autotrophically growing algae. Without an external carbon source, the relative activity of respiration is lower than photosynthesis and the impact of iron limitation on respiration appears minimal. In heterotrophic growth, however, iron limitation has a pronounced impact on respiration. Iron limitation has been shown in the literature to decrease respiration activity in heterotrophic bacteria [47, 50], yeast [9, 51, 52], and even protozoa [53].

It is hypothesized that in work presented here, low iron present in TGP cultures limits not only photosynthetic activity, but respiration as well. The complete oxidation of glucose to CO_2_ through respiration requires an investment of approximately 50 atoms of iron and produces about 30 ATP per glucose molecule, while the fermentation of glucose to lactate does not require any iron and produces 2 ATP per molecule of glucose [54]. When iron is limited, and glucose is available in abundance, *C. zofingiensis* is able to use fermentation pathways to supplement the energy loss that comes from lower TCA cycle activity. This behavior has been directly observed in *E. coli* during an experiment where cultures grown on glucose were subjected to increasing levels of iron limitation stress, which resulted in increasing levels of lactate excretion [54]. This has also been observed in yeast, when a fermentative substrate is provided in low iron conditions, the organism redirects its metabolism towards fermentation rather than respiration to prioritize iron use towards other necessary cell processes [9]. This would explain the observation presented here that that in *C. zofingiensis*, heterotrophic cultures grown on TGP media have high rates of glucose consumption and lactate secretion while heterotrophic cultures grown on nutrient dense CORE media do not.

### Conclusions

*Chromochloris zofingiensis* is capable of drastic changes to its metabolic phenotype. This green alga can switch from phototrophic growth to purely heterotrophic growth depending on nutrient conditions. During heterotrophic growth in low iron conditions, *C. zofingiensis* has been observed to ferment glucose to lactic acid even while in aerobic conditions. Iron is an essential nutrient for a number of cellular processes and iron limitation can have a pronounced impact on photosynthetic and respiratory activity.

In this work, we used isotopically assisted metabolic flux analysis to investigate the intracellular flux distributions associated with different phenotypes observed in heterotrophic and mixotrophic growth of *C. zofingiensis.* In cultures grown on TGP, a medium with low iron concentration, photosynthetic activity is shut down and the cell acts purely as a heterotroph even while under continuous illumination. In cultures grown on CORE media, a medium with high iron, with glucose under continuous illumination, the cells retain their photosynthetic capacity and display true mixotrophic growth. Additionally, TGP dark cultures display overflow metabolism, producing fermentation products while oxygen is available. Heterotrophic cultures grown in nutrient dense media do not ferment, and have full activity of the TCA cycle. This work provides further evidence of how metabolic function in *C. zofingiensis* can shift to acclimate to nutrient conditions and highlights the unique nature of this algae species. This rearrangement of metabolism to compensate for nutrient limitations has profound impacts on culture growth and productivity. Endeavors to compare productivities of an organism between different studies must account for the impact of different media formulations.

## Acknowledgments

Supported in part by US Department of Energy (DOE), Office of Biological and Environmental Research (BER) DE-SC0018301 and DE-SC0023027 to NB and SSM. The work on Fe poor cells was supported by the DOE grant DE-SC0020627 to SSM. DJC was supported in part by a training grant from the National Institutes of Health (NIH): 5T32GM007232-44.

A portion of this research was performed under the Facilities Integrating Collaborations for User Science (FICUS) program (https://doi.org/10.46936/fics.proj.2017.49960/60000021) and used resources at the DOE Joint Genome Institute (https://ror.org/04xm1d337) and the Environmental Molecular Sciences Laboratory (https://ror.org/04rc0xn13), which are DOE Office of Science User Facilities operated under Contract Nos. DE-AC02-05CH11231 (JGI) and DE-AC05-76RL01830 (EMSL).

