## Supplemental Information for "Investigating Overflow Metabolism in Heterotrophic Cultures of the Green Alga *Chromochloris zofingiensis*"

Supplemental Tables and Figures

**Supplemental Table S1**. List of metabolite abbreviations used.

| Abbreviation | Metabolite Name |
| --- | --- |
| GLC | Glucose |
| G6P | Glucose 6-phosphate |
| F6P | Fructose 6-phosphate |
| FBP | Fructose 1,6-bisphosphate |
| T3P | Triose 3-phosphate |
| GAP | Glyceraldehyde 3-phosphate |
| DHAP | Dihydroxyacetone phosphate |
| 13DHP | 1,3-Bisphosphoglycerate |
| 3PG | 3-phosphoglycerate |
| 2PG | 2-phosphoglycerate |
| PEP | Phosphoenolpyruvate |
| PYR | Pyruvate |
| ACCOA | Acetyl-CoA |
| CIT | Citrate |
| ICIT | Isocitrate |
| AKG | Alpha-ketoglutarate |
| SUC | Succinate |
| FUM | Fumarate |
| MAL | Malate |
| OA | Oxaloacetate |
| GLX | Glyoxylate |
| LAC | Lactate |
| P5P | Pentose 5-phosphate |
| R5P | Ribose 5-phosphate |
| X5P | Xylulose 5-phosphate |
| Ru5P | Ribulose 5-phosphate |
| Ru15BP | Ribulose 1,5-bisphosphate |
| S7P | Sedoheptulose 7-phosphate |
| S17BP | Sedoheptulose 1,7-bisphosphate |
| E4P | Erythrose 4-phosphate |
| CHOR | Chorismate |
| Phe | Phenylalanine |
| Tyr | Tyrosine |
| Leu | Leucine |
| Ser | Serine |
| Val | Valine |
| Ala | Alanine |
| Gly | Glycine |
| Glu | Glutamate |
| Asp | Aspartate |
| Thr | Threonine |

**Supplemental Table S2**. Metabolite fragments analyzed with mass spectrometry. Mass isotopomer distributions calculated for these metabolites were used in INCA to model intracellular fluxes. Cellular compartments are denoted as c (cytosol), h (chloroplast), and m (mitochondria).

| Metabolite | m/z | C atoms in fragment | Cellular Compartment |
| --- | --- | --- | --- |
| Alanine | 232 | 2 3 | c, m |
| Alanine | 260 | 1 2 3 | c, m |
| Glycine | 218 | 2 | m |
| Glycine | 246 | 1 2 | m |
| Valine | 260 | 2 3 4 5 | m |
| Valine | 288 | 1 2 3 4 5 | m |
| Leucine | 274 | 2 3 4 5 6 | h |
| Serine | 288 | 2 3 | h |
| Serine | 362 | 2 3 | h |
| Serine | 390 | 1 2 3 | h |
| Threonine | 376 | 2 3 4 | c |
| Threonine | 404 | 1 2 3 4 | c |
| Phenylalanine | 234 | 2 3 4 5 6 7 8 9 | h |
| Phenylalanine | 302 | 1 2 | h |
| Phenylalanine | 308 | 2 3 4 5 6 7 8 9 | h |
| Phenylalanine | 336 | 1 2 3 4 5 6 7 8 9 | h |
| Aspartate | 302 | 1 2 | c |
| Aspartate | 376 | 1 2 | c |
| Aspartate | 390 | 2 3 4 | c |
| Aspartate | 418 | 1 2 3 4 | c |
| Glutamate | 330 | 2 3 4 5 | m |
| Glutamate | 404 | 2 3 4 5 | m |
| Glutamate | 432 | 1 2 3 4 5 | m |
| Tyrosine | 302 | 1 2 | h |
| Alpha-ketoglutarate | 346 | 1 2 3 4 5 | m |
| Citrate | 459 | 1 2 3 4 5 6 | m |
| Fumarate | 287 | 1 2 3 4 | m |
| Lactate | 233 | 2 3 | m |
| Lactate | 261 | 1 2 3 | m |
| Malate | 419 | 1 2 3 4 | c, h, m |
| Pyruvate | 174 | 1 2 3 | c, h, m |
| Succinate | 289 | 1 2 3 4 | m |
| Arabinose | 284 | 1 2 3 4 | c |
| Galactose | 370 | 1 2 3 4 5 | c |
| Glucose (hydrolyzed starch) | 370 | 1 2 3 4 5 | h |
| Palmitic acid (16:0) | 74 | 1 2 | h |
| Palmitic acid (16:0) | 270 | 1 2 3 4 5 6 7 8 9 10 11 12 13 14 15 16 | h |
| Steric acid (18:0) | 74 | 1 2 | h |
| Steric acid (18:0) | 298 | 1 2 3 4 5 6 7 8 9 10 11 12 13 14 15 16 17 18 | h |
| Oleic acid (18:1) | 74 | 1 2 | h |
| Oleic acid (18:1) | 264 | 1 2 3 4 5 6 7 8 9 10 11 12 13 14 15 16 17 18 | h |
| Linoleic acid (18:2) | 74 | 1 2 | h |
| Linoleic acid (18:2) | 294 | 1 2 3 4 5 6 7 8 9 10 11 12 13 14 15 16 17 18 | h |

**Supplemental Table S3.** Complete list of reactions included in central metabolic network model of *C. zofingiensis.* Cellular compartments are denoted as E (external), c (cytosol), h (chloroplast), and m (mitochondria).

| Number | Reaction | Compartment | Pathway |
| --- | --- | --- | --- |
| 1 | Glc.e (abcdef) -> G6P.c (abcdef) | E | Exchange |
| 2 | G6P.c (abcdef) <-> G6P.h (abcdef) | c/h | Transport |
| 3 | G6P.h (abcdef) -> Ru5P.h (eadbc) + CO2.h (f) | h | Pentose Phosphate Pathway |
| 4 | Ru5P.h (abcde) <-> X5P.h (abcde) | h | Pentose Phosphate Pathway/ Calvin Cycle |
| 5 | Ru5P.h (abcde) <-> R5P.h (eadbc) | h | Pentose Phosphate Pathway/ Calvin Cycle |
| 6 | S7P.h (abcdefg) + T3P.h (hij) <-> R5P.h (bdfge) + X5P.h (aicjh) | h | Pentose Phosphate Pathway/ Calvin Cycle |
| 7 | S7P.h (abcdefg) + T3P.h (hij) <-> E4P.h (gbfd) + F6P.h (iajhec) | h | Pentose Phosphate Pathway |
| 8 | F6P.h (abcdef) + T3P.h (ghi) <-> E4P.h (eadc) + X5P.h (bhfig) | h | Pentose Phosphate Pathway |
| 9 | G6P.h (abcdef) <-> F6P.h (afbcde) | h | Glycolysis |
| 10 | F6P.h (abcdef) <-> T3P.h (ebf) + T3P.h (dac) | h | Glycolysis |
| 11 | Ru5P.h (abcde) -> Ru15bp.h (badce) | h | Calvin Cycle |
| 12 | Ru15bp.h (abcde) + CO2.h (f) -> 3PG.h (ace) + 3PG.h (bdf) | h | Calvin Cycle |
| 13 | T3P.h (abc) + E4P.h (defg) <-> S7P.h (becgafd) | h | Calvin Cycle |
| 14 | T3P.h (abc) <-> 3PG.h (bca) | h | Glycolysis / Calvin Cycle |
| 15 | T3P.h (abc) <-> T3P.c (abc) | c/h | Transport |
| 16 | T3P.c (abc) <-> PEP.c (bca) | c | Glycolysis |
| 17 | PEP.c (abc) <-> PYR.c (abc) | c | Glycolysis |
| 18 | PEP.c (abc) + CO2.c (d) -> OAA.c (abdc) | c | Pyruvate metabolism |
| 19 | PYR.c (abc) <-> PYR.m (abc) | c/m | Transport |
| 20 | PYR.m (abc) -> ACCoA.m (ab) + CO2.m (c) | m | Pyruvate metabolism |
| 21 | PYR.c (abc) -> PYR.h (abc) | c/h | Transport |
| 22 | PEP.h (abc) <-> PYR.h (abc) | h | Glycolysis |
| 23 | PYR.h (abc) -> ACCoA.h (ab) + CO2.h (c) | h | Pyruvate metabolism |
| 24 | ACCoA.m (ab) + OAA.m (cdef) -> CIT.m (caebfd) | m | TCA Cycle |
| 25 | CIT.m (abcdef) <-> ICIT.m (afcbed) | m | TCA Cycle |
| 26 | ICIT.m (abcdef) -> AKG.m (badcf) + CO2.m (e) | m | TCA Cycle |
| 27 | AKG.m (abcde) -> SUCC.m (badc) + CO2.m (e) | m | TCA Cycle |
| 28 | SUCC.m (abcd) -> FUM.m (abcd) | m | TCA Cycle |
| 29 | FUM.m (abcd) -> MAL.m (badc) | m | TCA Cycle |
| 30 | MAL.m (abcd) <-> OAA.m (abcd) | m | TCA Cycle |
| 31 | ICIT.m (abcdef) -> GLX.m (df) + SUCC.m (baec) | m | Glyoxylate shunt |
| 32 | ACCoA.m (ab) + GLX.m (cd) -> MAL.m (acbd) | m | Glyoxylate shunt |
| 33 | MAL.c (abcd) <-> OAA.c (abcd) | c | Redox/ Carbon shuttling |
| 34 | MAL.c (abcd) <-> MAL.m (abcd) | c/m | Transport |
| 35 | MAL.h (abcd) <-> OAA.h (abcd) | h | Redox/ Carbon shuttling |
| 36 | MAL.h (abcd) <-> PYR.h (abd) + CO2.h (c) | h | Redox/ Carbon shuttling |
| 37 | MAL.c (abcd) <-> PYR.c (abd) + CO2.c (c) | c | Redox/ Carbon shuttling |
| 38 | MAL.c (abcd) <-> MAL.h (abcd) | c/h | Transport |
| 39 | PYR.c (abc) -> LAC.c (abc) | c | Fermentation |
| 40 | AKG.m (abcde) -> Glu.b (abcde) | m | Amino acid synthesis |
| 41 | PYR.m (abc) -> Ala.b (abc) | m | Amino acid synthesis |
| 42 | PYR.c (abc) -> Ala.b (abc) | c | Amino acid synthesis |
| 43 | OAA.m (abcd) <-> Asp.m (abcd) | m | Redox/ Carbon shuttling |
| 44 | OAA.c (abcd) <-> Asp.c (abcd) | c | Redox/ Carbon shuttling |
| 45 | OAA.h (abcd) <-> Asp.h (abcd) | h | Redox/ Carbon shuttling |
| 46 | Asp.m (abcd) <-> Asp.c (abcd) | c/m | Transport |
| 47 | Asp.c (abcd) <-> Asp.h (abcd) | c/h | Transport |
| 48 | Asp.m (abcd) -> Asp.b (abcd) | m | Amino acid synthesis |
| 49 | Asp.c (abcd) -> Asp.b (abcd) | c | Amino acid synthesis |
| 50 | Asp.h (abcd) -> Asp.b (abcd) | h | Amino acid synthesis |
| 51 | Asp.h (abcd) -> Thr.b (cabd) | h | Amino acid synthesis |
| 52 | 3PG.h (abc) -> Ser.h (abc) | h | Amino acid synthesis |
| 53 | Ser.h (abc) -> PYR.h (abc) | h | Serine metabolism |
| 54 | Ser.h (abc) -> Ser.b (abc) | h | Amino acid synthesis |
| 55 | Ser.h (abc) -> Gly.h (bc) + thf.h (a) | h | Amino acid synthesis |
| 56 | GLX.m (ab) -> Gly.m (ab) | m | Amino acid synthesis |
| 57 | Gly.h (ab) -> Gly.b (ab) | h | Amino acid synthesis |
| 58 | Gly.m (ab) -> Gly.b (ab) | m | Amino acid synthesis |
| 59 | PEP.h (abc) + PEP.h (def) + E4P.h (ghij) -> chor.h (ahgibdjfce) | h | Amino acid synthesis |
| 60 | chor.h (abcdefghij) -> Tyr.b (bdchafgei) + CO2.h (j) | h | Amino acid synthesis |
| 61 | chor.h (abcdefghij) -> Phe.b (gchbdafei) + CO2.h (j) | h | Amino acid synthesis |
| 62 | PYR.h (abc) + PYR.h (def) -> Val.b (daebc) + CO2.h (f) | h | Amino acid synthesis |
| 63 | PYR.h (abc) + PYR.h (def) + ACCoA.h (gh) -> Leu.b (adbech) + CO2.h (g) + CO2.h (f) | h | Amino acid synthesis |
| 64 | G6P.c (abcdef) -> Ara.b (dfbce) + CO2.c (a) | c | Cell wall synthesis |
| 65 | G6P.c (abcdef) -> Gal.b (abcdef) | c | Cell wall synthesis |
| 66 | G6P.h (abcdef) <-> Sta.b (abcdef) | h | Starch synthesis |
| 67 | ACCoA.h (ab) + ACCoA.h (cd) + ACCoA.h (ef) + ACCoA.h (gh) + ACCoA.h (ij) + ACCoA.h (kl) + ACCoA.h (mn) + ACCoA.h (op) -> Pal.b (abcdefghijklmnop) | h | Fatty acid synthesis |
| 68 | ACCoA.h (ab) + ACCoA.h (cd) + ACCoA.h (ef) + ACCoA.h (gh) + ACCoA.h (ij) + ACCoA.h (kl) + ACCoA.h (mn) + ACCoA.h (op) + ACCoA.h (qr) -> Ster.b (abcdefghijklmnopqr) | h | Fatty acid synthesis |
| 69 | ACCoA.h (ab) + ACCoA.h (cd) + ACCoA.h (ef) + ACCoA.h (gh) + ACCoA.h (ij) + ACCoA.h (kl) + ACCoA.h (mn) + ACCoA.h (op) + ACCoA.h (qr) -> Ole.b (abcdefghijklmnopqr) | h | Fatty acid synthesis |
| 70 | ACCoA.h (ab) + ACCoA.h (cd) + ACCoA.h (ef) + ACCoA.h (gh) + ACCoA.h (ij) + ACCoA.h (kl) + ACCoA.h (mn) + ACCoA.h (op) + ACCoA.h (qr) -> Lino.b (abcdefghijklmnopqr) | h | Fatty acid synthesis |
| 71 | CO2.e (a) -> CO2.c (a) | E | Exchange |
| 72 | CO2.c (a) <-> CO2.h (a) | c/h | Transport |
| 73 | CO2.c (a) <-> CO2.m (a) | c/m | Transport |
| 74 | LAC.c (abc) -> LAC.e (abc) | E | Exchange |
| 75 | SUCC.m (abcd) -> SUCC.e (abcd) | E | Exchange |

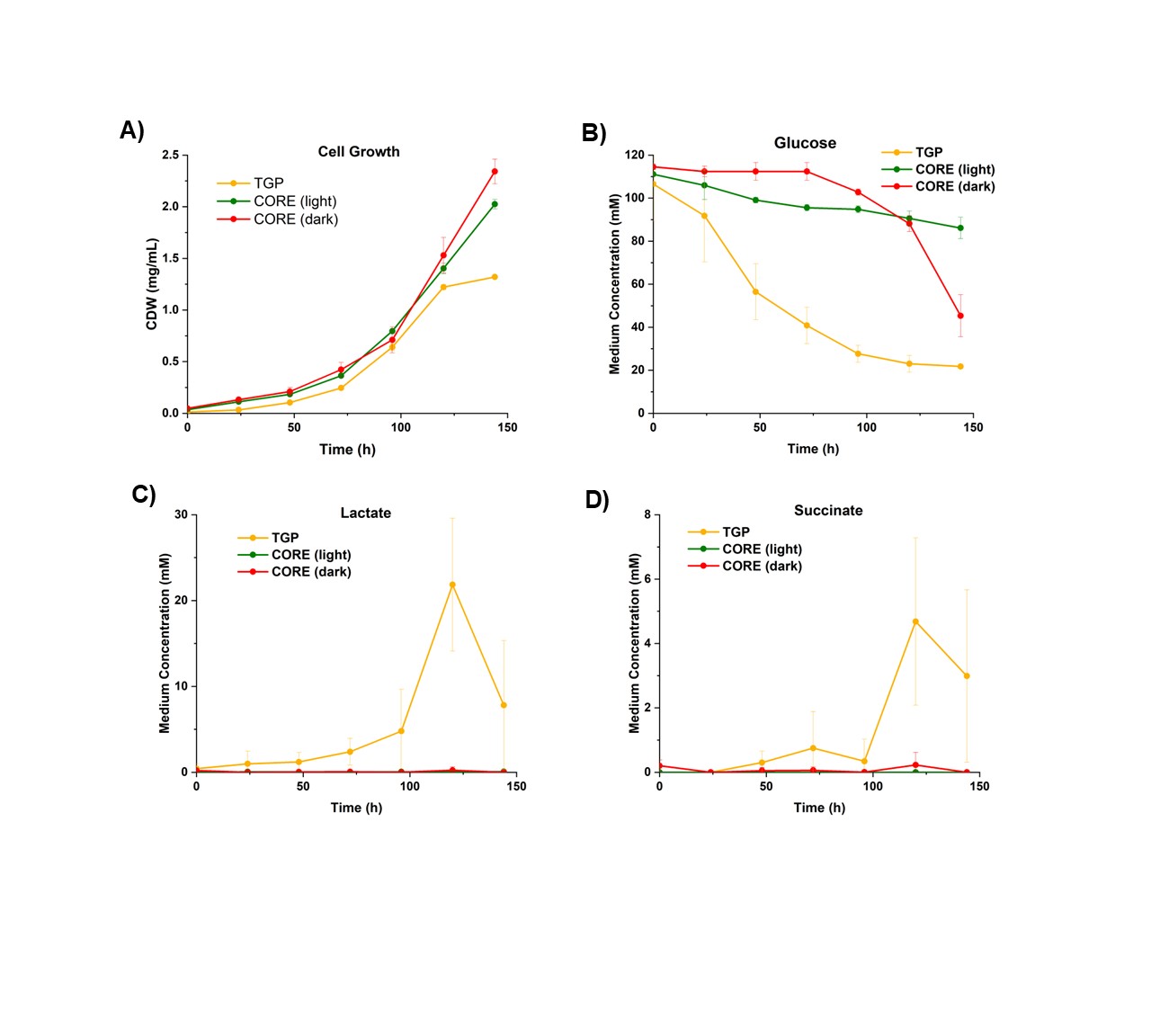

**Supplemental Figure S1**. A) Cell dry weight data, B) glucose consumption data, C) lactate excretion data, and D) succinate excretion data over time for *C. zofingiensis* cultures grown in TGP medium in continuous light, CORE medium in continuous light, and CORE medium in the dark.

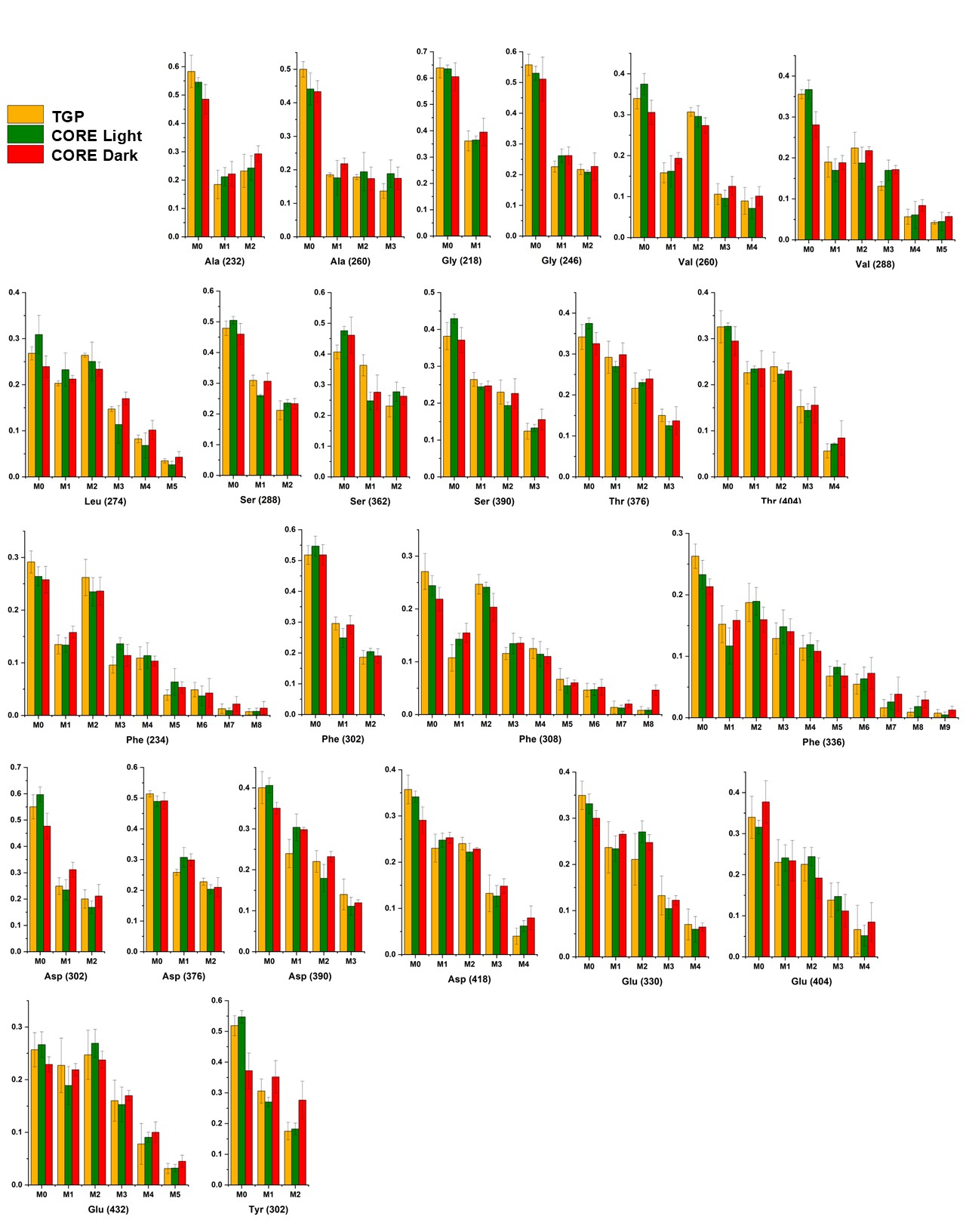

**Supplemental Figure S2**. Mass isotopomer distributions for amino acids. Amino acid fragment m/z values provided in parenthesis. Error bars represent standard deviation of n=6 measurements.

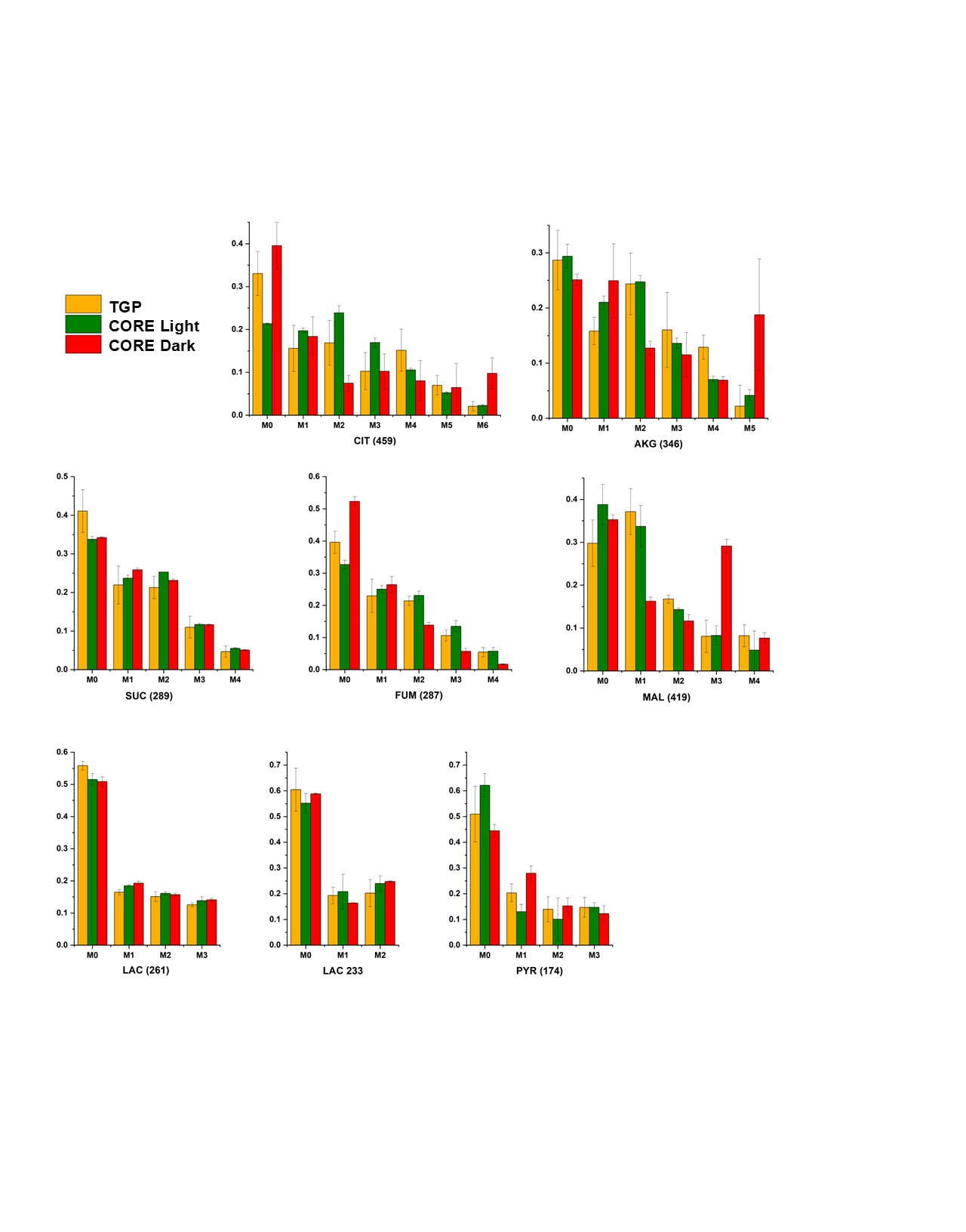

**Supplemental Figure S3**. Mass isotopomer fragments for organic acids. Fragment m/z values are provided in parenthesis. Error bars represent standard deviation of n=3 measurements.

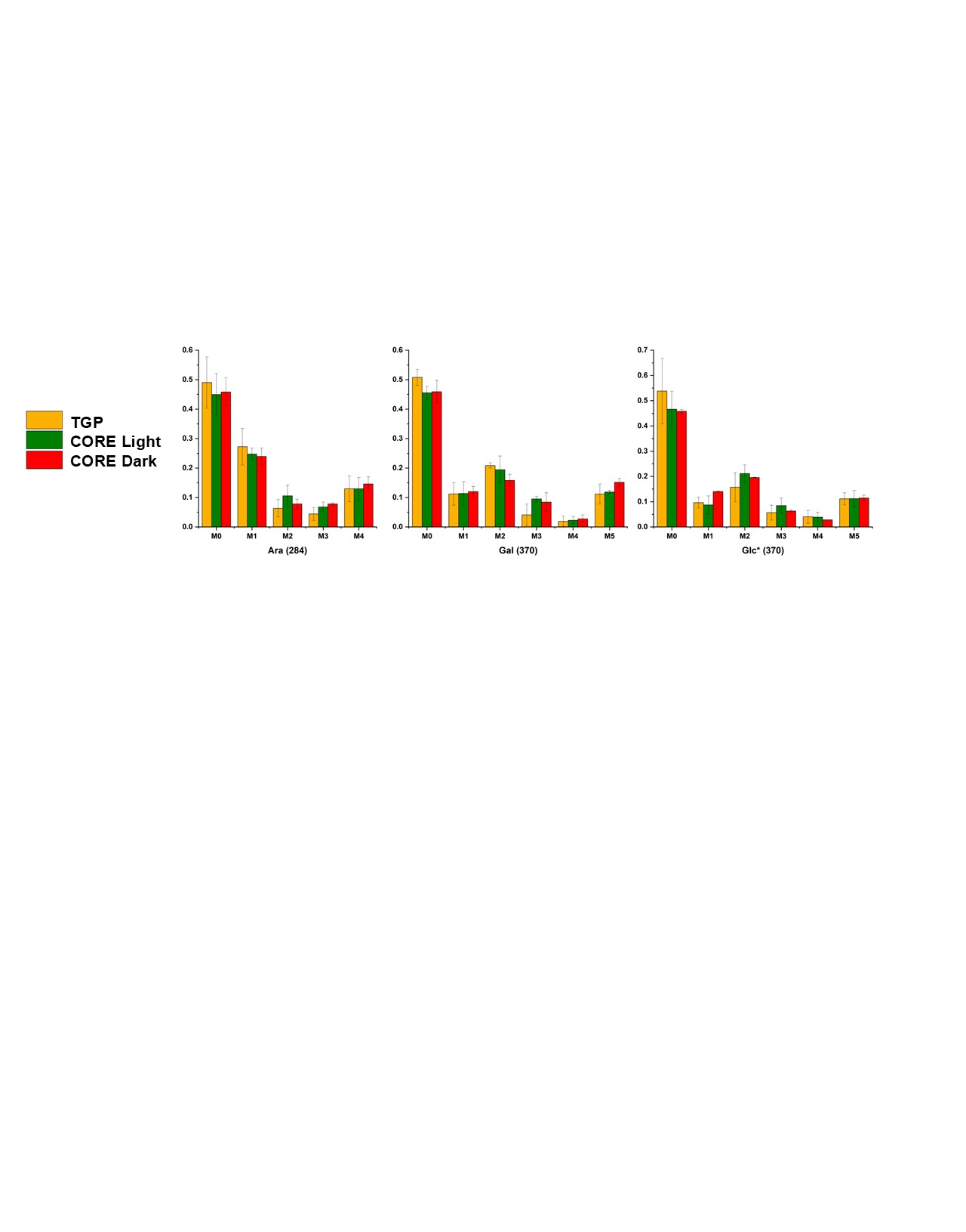

**Supplemental Figure S4**. Mass isotopomer fragments for sugars. Fragment m/z values are provided in parenthesis. Error bars represent standard deviation of n=3 measurements. Asterisk denotes isotopomer abundances from glucose derived from starch.

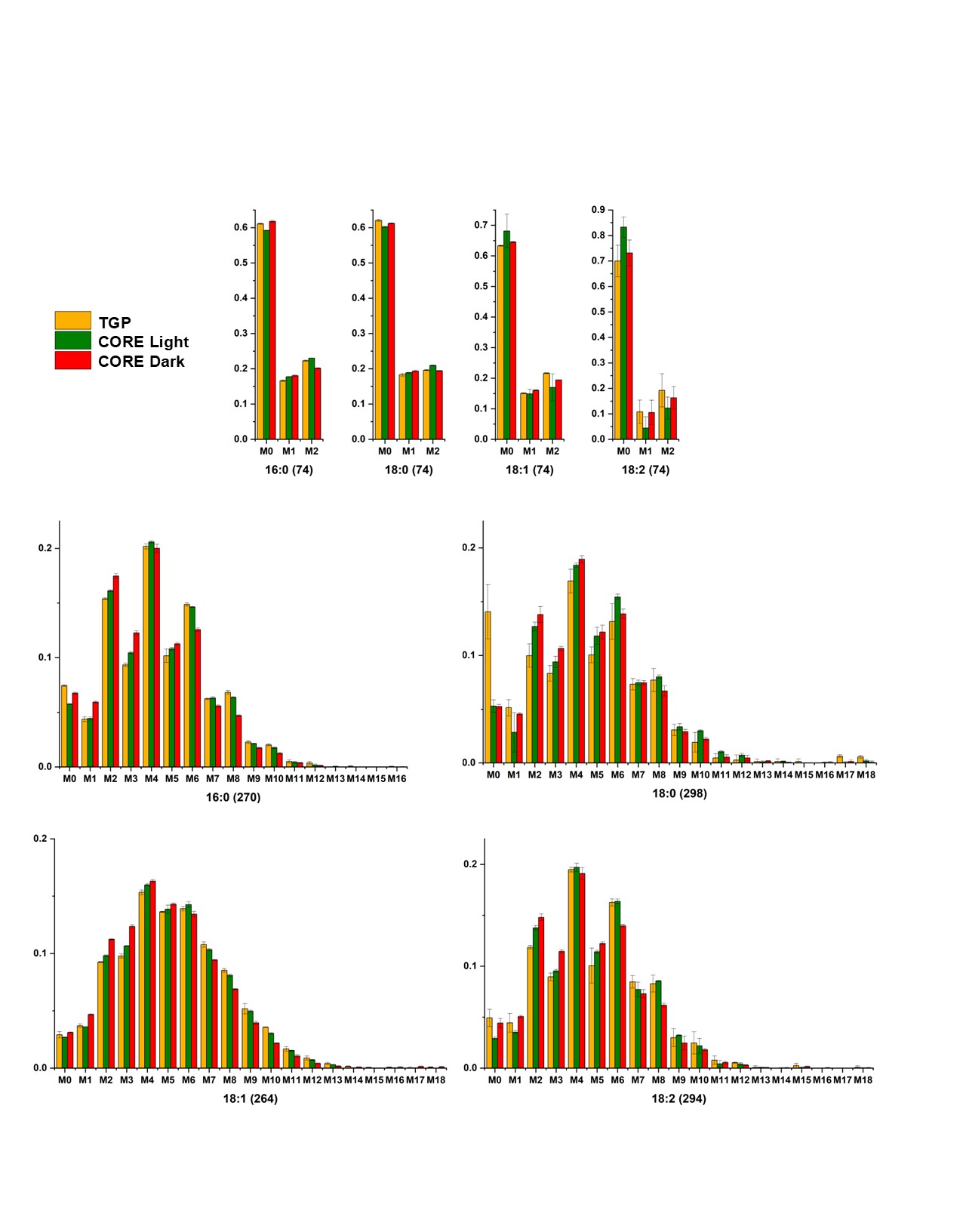

**Supplemental Figure S5**. Mass isotopomer fragments for fatty acids. Fragment m/z values are provided in parenthesis. Error bars represent standard deviation of n=3 measurements.

**Supplemental Table S4**. Mean FPKMs and log_2_ fold change values for transcriptomics data collected for genes associated with mitochondrial TCA cycle and glyoxylate shunt reactions in low and very low iron conditions.

|  |  | Mean FPKM | | | Log_2_ Fold Change Compared to Replete | |
| --- | --- | --- | --- | --- | --- | --- |
| Reaction catalyzed by associated enzyme | Gene ID | Iron Replete (20 µM) | Low Fe (2 µM) | Very Low Fe (0.2 µM) | Low Fe (2 µM) | Very Low Fe (0.2 µM) |
| PYR -> CO2 + ACCOA | Cz08g00230 | 67.6 | 56.4 | 66.3 | -0.3 | 0.0 |
| PYR -> CO2 + ACCOA | Cz03g08090 | 372.7 | 202.9 | 238.1 | -0.9 | -0.6 |
| PYR -> CO2 + ACCOA | Cz18g13050 | 145.2 | 128.9 | 125.7 | -0.2 | -0.2 |
| ACCOA + OA -> CIT | Cz02g12210 | 145.4 | 152.4 | 161.9 | 0.1 | 0.1 |
| ACCOA + OA -> CIT | Cz02g27080 | 46.8 | 47.9 | 43.9 | 0.0 | -0.1 |
| ACCOA + OA -> CIT | Cz06g12020 | 66.4 | 41.2 | 36.8 | -0.7 | -0.9 |
| CIT <-> ICIT | Cz13g00140 | 133.7 | 97.3 | 121.3 | -0.5 | -0.1 |
| ICIT -> AKG + CO2 | Cz12g16160 | 162.2 | 127.7 | 129.0 | -0.3 | -0.3 |
| ICIT -> AKG + CO2 | Cz11g28180 | 100.7 | 116.1 | 116.7 | 0.2 | 0.2 |
| ICIT -> AKG + CO2 | Cz11g08120 | 103.6 | 85.2 | 89.5 | -0.3 | -0.2 |
| ICIT -> AKG + CO2 | Cz12g16160 | 162.2 | 127.7 | 129.0 | -0.3 | -0.3 |
| AKG -> SUC + CO2 | Cz05g03220 | 146.2 | 117.9 | 135.1 | -0.3 | -0.1 |
| AKG -> SUC + CO2 | Cz01g28100 | 29.7 | 22.7 | 21.9 | -0.4 | -0.4 |
| AKG -> SUC + CO2 | Cz02g11010 | 62.7 | 72.6 | 87.0 | 0.2 | 0.5 |
| AKG -> SUC + CO2 | Cz01g33150 | 92.5 | 86.7 | 106.4 | -0.1 | 0.2 |
| SUC <-> FUM | Cz07g14020 | 85.5 | 117.1 | 110.3 | 0.4 | 0.4 |
| SUC <-> FUM | Cz15g13230 | 68.7 | 72.5 | 79.9 | 0.1 | 0.2 |
| FUM <-> MAL | Cz07g04030 | 98.6 | 93.8 | 79.6 | -0.1 | -0.3 |
| MAL <-> OA | Cz04g33150 | 64.9 | 51.2 | 55.9 | -0.3 | -0.2 |
| MAL <-> OA | Cz01g16230 | 209.1 | 141.4 | 129.2 | -0.6 | -0.7 |
| ICIT -> GLX + SUC | Cz07g04220 | 0.8 | 0.5 | 1.2 | -0.3 | 0.4 |
| GLX + ACCOA -> MAL | Cz14g25060 | 7.4 | 20.0 | 21.8 | 1.4 | 1.5 |

**Supplemental Table S5**. Mean FPKMs and log_2_ fold change values for transcriptomics data collected for genes associated with chloroplast CBB cycle and PPP reactions in low and very low iron conditions.

|  |  | Mean FPKM | | | Log_2_ Fold Change Compared to Replete | |
| --- | --- | --- | --- | --- | --- | --- |
| Reaction catalyzed by associated enzyme | Gene ID | Iron Replete (20 µM) | Low Fe (2 µM) | Very Low Fe (0.2 µM) | Low Fe (2 µM) | Very Low Fe (0.2 µM) |
| Ru5P -> RU15BP | Cz07g08090 | 698.4 | 626.8 | 786.0 | -0.1 | 0.2 |
| Ru15BP + CO2 -> 2*3PG | Cz17g13100 | 15103.8 | 7559.3 | 9628.1 | -1.0 | -0.6 |
| Ru15BP + CO2 -> 2*3PG | CzCPg00360 | 9.0 | 1.8 | 0.9 | -2.1 | -3.2 |
| 3PG -> 13DPG | Cz16g19260 | 512.7 | 286.7 | 374.3 | -0.8 | -0.4 |
| 13DPG <-> GAP | Cz05g34160 | 502.5 | 477.2 | 643.1 | -0.1 | 0.3 |
| 13DPG <-> GAP | Cz11g09290 | 1692.0 | 727.7 | 826.9 | -1.2 | -1.0 |
| GAP <-> DHAP | Cz06g17270 | 171.8 | 84.7 | 83.1 | -1.0 | -1.0 |
| DHAP + GAP -> FBP | Cz03g06050 | 70.7 | 21.4 | 25.4 | -1.7 | -1.5 |
| DHAP + GAP -> FBP | Cz05g37140 | 4704.1 | 2275.7 | 2778.2 | -1.0 | -0.8 |
| DHAP + GAP -> FBP | Cz06g07090 | 45.9 | 43.1 | 43.6 | -0.1 | -0.1 |
| DHAP + GAP -> FBP | Cz03g37060 | 30.2 | 30.9 | 32.3 | 0.0 | 0.1 |
| DHAP + GAP -> FBP | Cz04g03070 | 156.5 | 86.2 | 84.7 | -0.8 | -0.9 |
| FBP -> F6P | Cz03g37060 | 30.2 | 30.9 | 32.3 | 0.0 | 0.1 |
| GAP + F6P <-> E4P + F6P | Cz04g12210 | 90.8 | 75.2 | 60.1 | -0.3 | -0.6 |
| E4P + DHAP -> S17BP | Cz06g07090 | 45.9 | 43.1 | 43.6 | -0.1 | -0.1 |
| S17BP -> S7P | Cz01g42020 | 523.1 | 215.2 | 241.5 | -1.3 | -1.1 |
| X5P + R5P <-> G3P + S7P | Cz03g04080 | 573.7 | 479.9 | 592.9 | -0.2 | 0.0 |
| X5P <-> Ru5P | Cz05g11190 | 183.3 | 134.9 | 154.2 | -0.4 | -0.3 |
| X5P <-> Ru5P | Cz14g07140 | 5.7 | 6.1 | 7.0 | 0.1 | 0.3 |
| R5P <-> Ru5P | Cz09g17220 | 191.7 | 64.4 | 57.4 | -1.6 | -1.7 |
| R5P <-> Ru5P | Cz05g13260 | 18.5 | 18.7 | 19.2 | 0.0 | 0.0 |

**Supplemental Table S6**. Mean FPKMs and log_2_ fold change values for transcriptomics data collected for genes associated with mitochondrial respiration electron transport chain in low and very low iron conditions.

|  |  | Mean FPKM | | | Log_2_ Fold Change Compared to Replete | |
| --- | --- | --- | --- | --- | --- | --- |
| Associated Protein | Gene ID | Iron Replete (20 µM) | Low Fe (2 µM) | Very Low Fe (0.2 µM) | Low Fe (2 µM) | Very Low Fe (0.2 µM) |
| Complex I | CzMTg00150 | 2.0 | 1.3 | 1.0 | -0.3 | -0.6 |
| Complex I | Cz16g19250 | 162.8 | 141.4 | 146.4 | -0.2 | -0.2 |
| Complex I | Cz14g19020 | 66.4 | 65.0 | 70.6 | 0.0 | 0.1 |
| Complex I | Cz12g16270 | 22.6 | 32.4 | 30.4 | 0.5 | 0.4 |
| Complex I | Cz12g00190 | 83.3 | 82.7 | 94.3 | 0.0 | 0.2 |
| Complex I | Cz09g05210 | 137.5 | 119.2 | 121.3 | -0.2 | -0.2 |
| Complex I | Cz05g18120 | 44.9 | 40.7 | 45.7 | -0.1 | 0.0 |
| Complex I | Cz02g06240 | 200.1 | 215.8 | 189.8 | 0.1 | -0.1 |
| Complex II | Cz05g08055 | 14.3 | 27.4 | 25.2 | 0.9 | 0.8 |
| Complex II | Cz15g13230 | 68.7 | 72.5 | 79.9 | 0.1 | 0.2 |
| Complex III | Cz13g13110 | 305.8 | 237.6 | 222.9 | -0.4 | -0.5 |
| Complex III | Cz10g27170 | 249.3 | 225.4 | 220.9 | -0.1 | -0.2 |
| Complex III | Cz05g15120 | 295.1 | 112.3 | 102.5 | -1.4 | -1.5 |
| Complex IV | Cz03g08180 | 263.6 | 193.9 | 229.5 | -0.4 | -0.2 |
| Complex IV | Cz01g24170 | 246.1 | 202.4 | 244.4 | -0.3 | 0.0 |
| Complex IV | Cz04g33210 | 293.4 | 353.8 | 283.2 | 0.3 | 0.0 |
| Complex IV | Cz05g15120 | 295.1 | 112.3 | 102.5 | -1.4 | -1.5 |
| Complex IV | Cz11g20250 | 248.3 | 231.8 | 197.3 | -0.1 | -0.3 |
| Complex IV | Cz15g01180 | 176.6 | 210.7 | 179.6 | 0.3 | 0.0 |
| Complex IV | CzMTg00040 | 2.6 | 1.2 | 1.6 | -0.9 | -0.6 |
| Complex IV | CzMTg00100 | 3.3 | 3.0 | 2.6 | -0.1 | -0.4 |
| ATP Synthase | Cz04g31080 | 279.0 | 239.0 | 205.1 | -0.2 | -0.4 |
| ATP Synthase | Cz04g32030 | 381.8 | 314.4 | 254.7 | -0.3 | -0.6 |
| ATP Synthase | Cz05g16180 | 196.5 | 159.7 | 162.6 | -0.3 | -0.3 |
| ATP Synthase | Cz08g05010 | 330.8 | 280.5 | 282.5 | -0.2 | -0.2 |
| ATP Synthase | Cz09g21130 | 583.3 | 345.2 | 382.9 | -0.7 | -0.6 |
| ATP Synthase | Cz11g25250 | 113.5 | 88.8 | 85.7 | -0.3 | -0.4 |
| ATP Synthase | Cz12g25090 | 13.7 | 10.6 | 11.6 | -0.4 | -0.2 |
| ATP Synthase | CzMTg00140 | 54.3 | 33.2 | 45.4 | -0.6 | -0.2 |
| ATP Synthase | Cz03g10280 | 13.1 | 11.6 | 13.1 | -0.2 | 0.0 |
| ATP Synthase | Cz03g14200 | 351.0 | 341.4 | 321.1 | 0.0 | -0.1 |
| ATP Synthase | Cz03g32200 | 488.8 | 398.8 | 368.0 | -0.3 | -0.4 |
| ATP Synthase | Cz04g09240 | 255.8 | 208.2 | 206.3 | -0.3 | -0.3 |
| ATP Synthase | Cz04g32030 | 381.8 | 314.4 | 254.7 | -0.3 | -0.6 |
| ATP Synthase | Cz05g29070 | 270.5 | 257.7 | 219.4 | -0.1 | -0.3 |
| ATP Synthase | Cz06g10150 | 236.0 | 207.3 | 191.5 | -0.2 | -0.3 |
| ATP Synthase | Cz06g20230 | 23.6 | 21.1 | 21.5 | -0.2 | -0.1 |
| ATP Synthase | Cz06g37200 | 443.1 | 399.3 | 338.8 | -0.1 | -0.4 |
| ATP Synthase | Cz08g05010 | 330.8 | 280.5 | 282.5 | -0.2 | -0.2 |
| ATP Synthase | Cz18g09050 | 164.9 | 149.4 | 142.9 | -0.1 | -0.2 |
| ATP Synthase | CzMTg00230 | 3.2 | 2.3 | 1.8 | -0.4 | -0.8 |

**Supplemental Table S7**. Mean FPKMs and log_2_ fold change values for transcriptomics data collected for genes associated with chloroplast photosynthetic electron transport chain in low and very low iron conditions.

|  |  | Mean FPKM | | | Log_2_ Fold Change Compared to Replete | |
| --- | --- | --- | --- | --- | --- | --- |
| Associated Protein | Gene ID | Iron Replete (20 µM) | Low Fe (2 µM) | Very Low Fe (0.2 µM) | Low Fe (2 µM) | Very Low Fe (0.2 µM) |
| PSI | Cz01g23030 | 521.0 | 398.5 | 575.9 | -0.3 | 0.1 |
| PSI | Cz02g14140 | 387.9 | 520.2 | 705.5 | 0.4 | 0.8 |
| PSI | Cz04g04130 | 532.6 | 605.4 | 791.6 | 0.2 | 0.5 |
| PSI | Cz04g09210 | 724.9 | 260.5 | 235.0 | -1.4 | -1.6 |
| PSI | Cz07g02100 | 2031.7 | 311.3 | 583.7 | -2.7 | -1.8 |
| PSI | Cz07g28150 | 1811.4 | 624.8 | 829.5 | -1.5 | -1.1 |
| PSI | Cz09g32070 | 769.3 | 357.2 | 453.6 | -1.0 | -0.7 |
| PSI | Cz10g10160 | 911.5 | 1185.7 | 1417.3 | 0.4 | 0.6 |
| PSI | Cz11g23240 | 14.2 | 13.5 | 15.7 | -0.1 | 0.1 |
| PSI | Cz12g16070 | 573.3 | 151.3 | 153.5 | -1.8 | -1.8 |
| PSI | Cz13g08120 | 266.9 | 248.9 | 277.0 | -0.1 | 0.0 |
| PSI | Cz14g01080 | 325.8 | 2280.1 | 2781.8 | 2.7 | 3.0 |
| PSI | Cz15g17110 | 828.2 | 144.3 | 180.9 | -2.5 | -2.2 |
| PSI | Cz16g14150 | 849.0 | 197.2 | 195.8 | -2.1 | -2.1 |
| PSI | Cz01g05275 | 769.6 | 249.7 | 244.1 | -1.6 | -1.6 |
| PSI | Cz02g18180 | 1007.8 | 359.6 | 398.2 | -1.4 | -1.3 |
| PSI | Cz04g01180 | 863.2 | 417.9 | 276.5 | -1.0 | -1.6 |
| PSI | Cz05g36030 | 1196.2 | 344.6 | 392.2 | -1.8 | -1.6 |
| PSI | Cz06g26010 | 734.4 | 275.2 | 211.3 | -1.4 | -1.8 |
| PSI | Cz06g31280 | 314.4 | 98.2 | 91.1 | -1.6 | -1.8 |
| PSI | Cz13g06010 | 760.7 | 252.4 | 267.2 | -1.6 | -1.5 |
| PSI | Cz15g01010 | 725.7 | 290.0 | 247.1 | -1.3 | -1.5 |
| PSI | Cz17g10010 | 541.4 | 153.0 | 130.7 | -1.7 | -2.0 |
| PSI | Cz01g35130 | 1635.6 | 831.7 | 1265.3 | -0.9 | -0.4 |
| PSI | Cz02g37160 | 106.8 | 82.8 | 82.6 | -0.4 | -0.4 |
| PSI | Cz15g17110 | 828.2 | 144.3 | 180.9 | -2.5 | -2.2 |
| PSI | Cz07g02110 | 2142.5 | 3109.8 | 4300.8 | 0.5 | 1.0 |
| PSII | Cz01g04040 | 69.4 | 40.8 | 40.4 | -0.7 | -0.8 |
| PSII | Cz01g23030 | 521.0 | 398.5 | 575.9 | -0.3 | 0.1 |
| PSII | Cz02g14140 | 387.9 | 520.2 | 705.5 | 0.4 | 0.8 |
| PSII | Cz02g31140 | 22.9 | 24.8 | 23.1 | 0.1 | 0.0 |
| PSII | Cz03g01250 | 32.1 | 25.0 | 24.1 | -0.2 | -0.3 |
| PSII | Cz03g26240 | 980.3 | 384.0 | 354.5 | -1.3 | -1.4 |
| PSII | Cz03g38250 | 20.9 | 18.9 | 14.5 | -0.1 | -0.5 |
| PSII | Cz04g04130 | 532.6 | 605.4 | 791.6 | 0.2 | 0.5 |
| PSII | Cz04g24050 | 0.2 | 0.2 | 0.1 | 0.0 | -0.2 |
| PSII | Cz06g29040 | 1.4 | 0.8 | 0.9 | -0.7 | -0.6 |
| PSII | Cz07g02100 | 2031.7 | 311.3 | 583.7 | -2.7 | -1.8 |
| PSII | Cz07g02110 | 2142.5 | 3109.8 | 4300.8 | 0.5 | 1.0 |
| PSII | Cz08g11160 | 1097.3 | 220.3 | 225.2 | -2.3 | -2.2 |
| PSII | Cz11g23240 | 14.2 | 13.5 | 15.7 | -0.1 | 0.1 |
| PSII | Cz12g16070 | 573.3 | 151.3 | 153.5 | -1.8 | -1.8 |
| PSII | Cz13g03240 | 4180.7 | 587.7 | 356.5 | -2.8 | -3.5 |
| PSII | Cz15g09060 | 1893.2 | 175.8 | 234.4 | -3.4 | -3.0 |
| PSII | UNPLg00080 | 3095.6 | 694.6 | 469.2 | -2.1 | -2.7 |
| PSII | Cz02g18040 | 104.7 | 41.9 | 34.5 | -1.3 | -1.6 |
| PSII | Cz03g13190 | 28.0 | 17.7 | 27.3 | -0.6 | 0.0 |
| PSII | Cz04g12120 | 10.3 | 8.5 | 9.4 | -0.3 | -0.1 |
| PSII | Cz07g15040 | 982.0 | 256.5 | 260.8 | -1.9 | -1.9 |
| PSII | Cz09g21140 | 1169.4 | 1040.2 | 1183.7 | -0.2 | 0.0 |
| PSII | Cz10g21040 | 92.5 | 45.2 | 48.6 | -1.0 | -0.9 |
| PSII | Cz12g14170 | 47.8 | 29.1 | 41.4 | -0.7 | -0.2 |
| PSII | Cz16g19040 | 1413.2 | 471.7 | 426.2 | -1.5 | -1.7 |
| PSII | Cz19g02070 | 28.7 | 15.6 | 18.5 | -0.9 | -0.6 |
| PSII | Cz04g12110 | 150.5 | 63.7 | 58.3 | -1.2 | -1.4 |
| PSII | Cz05g09100 | 788.4 | 200.4 | 210.8 | -1.9 | -1.9 |
| PSII | Cz13g07140 | 84.2 | 43.8 | 52.9 | -0.9 | -0.7 |
| PSII | Cz17g15140 | 392.9 | 186.9 | 181.0 | -1.0 | -1.1 |
| cytochome b6/f | Cz06g16040 | 283.9 | 217.4 | 217.0 | -0.4 | -0.4 |
| cytochome b6/f | Cz16g12040 | 416.4 | 244.9 | 227.0 | -0.7 | -0.9 |
| cytochome b6/f | Cz01g35130 | 1635.6 | 831.7 | 1265.3 | -0.9 | -0.4 |
| cytochome b6/f | Cz13g12160 | 420.3 | 174.1 | 151.8 | -1.2 | -1.4 |
| ATP synthase | Cz02g27060 | 605.8 | 353.7 | 301.4 | -0.7 | -1.0 |
| ATP synthase | Cz09g21130 | 583.3 | 345.2 | 382.9 | -0.7 | -0.6 |
| ATP synthase | Cz08g05010 | 330.8 | 280.5 | 282.5 | -0.2 | -0.2 |
| ATP synthase | Cz05g34180 | 557.5 | 354.5 | 325.1 | -0.6 | -0.8 |
